# Bacterial endosymbionts influence fungal transcriptional profiles with implications for host response in the human fungal pathogens *Rhizopus microsporus* and *Rhizopus delemar*

**DOI:** 10.1101/580746

**Authors:** Poppy Sephton-Clark, José F Muñoz, Herbert Itabangi, Kerstin Voelz, Christina A Cuomo, Elizabeth R Ballou

## Abstract

Mucorales spores, the causative agents of mucormycosis, interact with the innate immune system to cause acute, chronic, or resolving infection. Understanding the factors that influence disease initiation and progression is key to understanding mucormycosis and developing new treatments. Complicating this, mucormycosis can be caused by a number of species that span the Mucorales order and may be host to bacterial endosymbionts. This study sets out to examine the differences between two species in the Mucorales order by characterising their differential interactions with the innate immune system, and their interactions with environmental bacterial endosymbionts. Through a holistic approach, this study examines the transcriptional responses of *Rhizopus delemar* and *Rhizopus microsporus*, two of the most commonly diagnosed species, to innate immune cells. This study also examines the immune cell response and assesses the variation in these responses, given the presence or absence of bacterial endosymbionts within the fungi. We see that the fungal response is driven by interaction with innate immune cells. Moreover, the effect of the bacterial endosymbiont on the fungus is species-specific and strongly influences fungal transcription during phagocyte stress. The macrophage response varies depending on the infecting fungal species, and depending on endosymbiont status. Macrophages are better able to survive when germination is inhibited, or upon a pro-inflammatory response. This work reveals species-specific host responses to related Mucorales spores and shows that bacterial endosymbionts have an important role to play by impacting both innate immune cell response, and fungal response when under stress.

## Introduction

Mucormycosis is a devastating, environmentally acquired fungal infection caused by a variety of Mucorales species. Understanding the interaction between Mucorales spores and innate immune cells is key to understanding mucormycosis. Studies frequently focus upon the interaction of a single species of the Mucorales order with innate immune cells (Warris *et al*. 2005; Chamilos *et al*. 2008; Schmidt *et al*. 2013; Kraibooj *et al*. 2014; Inglesfield *et al*. 2018). However, between Mucorales species, and across infecting isolates, numerous phenotypic and genomic differences can be observed (Hoffmann *et al*. 2013; Mendoza *et al*. 2014; Schwartze *et al*. 2014), and the range of virulence profiles displayed across the family indicates there may be diversified mechanisms at work (Petraitis *et al*. 2013). Pathogenicity of these species has been linked to the variable copy number of CotH, a family of cell surface proteins important for spores’ interactions with and adherence to endothelial cells. Iron-scavenging and melanisation pathways have also been shown to play a role in pathogenicity (Chibucos *et al*. 2016; Andrianaki *et al*. 2018). In addition to fungal genome plasticity and transcriptional responses to host stimuli, fungal-host interactions can be further modulated by bacterial endosymbionts. Endosymbionts have been shown to produce secondary metabolites that modulate pathogenesis, and can directly influence fungal transcription (Partida-Martinez & Hertweck 2005; Itabangi *et al*. 2019; Carter *et al*. 2020). The majority of Mucorales species harbour bacterial endosymbionts, and species can vary between isolates of the same fungal host (Ibrahim *et al*. 2008; Kobayashi & Crouch 2009; Mondo *et al*. 2017; Itabangi *et al*. 2019). For example, *R. microsporus* has been shown to harbour and be modulated by interaction with *Mycetohabitans endofungorum* and *Mycetohabitan rhizoxinica* (formerly *Burkholderia endofungorum* and *Burkholderia rhizoxinica)* (Partida-Martinez & Hertweck 2005; Estrada-de los Santos *et al*. 2018). We have recently shown that a bacterial endosymbiont, *Ralstonia pickettii,* influences the outcome of *Rhizopus microsporus* infections in both zebrafish and murine models through modulation of both fungal and phagocyte phenotypes (Itabangi *et al*. 2019). However, the impact of this endosymbiont on fungal and host transcriptional responses, and the relevance of endosymbiosis to fungal-mammalian host interactions across Mucorales species, remains uncharacterised.

Within the Mucorales order, *Rhizopus delemar* and *Rhizopus microsporus* are responsible for the majority of infections within immune compromised individuals (Liu *et al*. 2018). Risk factors for developing the disease include, but are not limited to: innate immune cell deficiencies, uncontrolled diabetes, immune suppression, skin barrier breaches and iron overload (Baldin and Ibrahim 2017). With limited antifungal treatments effective against these invasive infections, there is often a need for surgical debridement or amputation (Spellberg *et al*. 2005). In disseminated cases, mucormycosis may present with a mortality rate of over 90% (Mendoza *et al*. 2014). Reports of chronic mucormycosis have also emerged, affecting both immunocompromised and immunocompetent patients (Alan 2019).

The innate immune response to Mucorales spores is key to infection control: macrophages are required to induce a differential immune response on contact with *Rhizopus* spp. and, in healthy individuals, may inhibit germination or kill hyphal forms upon contact (Ghuman and Voelz 2017, Andrianaki *et al*. 2018). Host-mediated iron limitation allows macrophages to more effectively kill *Rhizopus* spp. (Ibrahim *et al*. 2008; Liu *et al*. 2015; Andrianaki *et al*. 2018). Single and multi-species studies have shown that phagosome maturation is arrested by melanin within the cell walls of *Aspergillus* spp. and *Rhizopus* spp., and several works comparing *Aspergillus* spp. to *Rhizopus* spp. have revealed similar immunostimulatory capacities, but differences in their responses to host stress (Warris *et al*. 2005; Chamilos *et al*. 2008; Schmidt *et al*. 2013; Kraibooj *et al*. 2014). Exploring and understanding fungal responses to the host is essential to improving our understanding of mucormycosis, yet it remains unclear how Mucorales species respond to, and interact with, the innate immune system, and to what extent this varies by species and by holo-genome.

Our work explores the interplay between the innate immune system, *R. delemar,* and *R. microsporus,* and their respective endosymbionts. We investigate the differences between these two fungal species, how they respond transcriptionally to innate immune cells, and how their respective bacterial endosymbionts affect this interaction. We also investigate the transcriptional response of innate immune cells to these infectious spores and determine how this interaction is influenced by the presence of an endosymbiont. We reveal a large difference between the fungal transcriptional profiles of *R. delemar* and *R. microsporus* during *in vitro* monoculture. There is a small conserved response on exposure to innate immune cells, including key changes in cell wall genes, consistent with germination. Conversely, we see that the immune cell response differs significantly between fungal species and is also influenced by the presence or lack of an endosymbiont. We also observe that through the activation of innate immune cells, or upon inhibition of chitin synthase, we can improve the ability of innate immune cells to control fungal spores. Our work represents a broad analysis of the transcriptional interplay between innate immune responders, infectious Mucorales spores, and their respective endosymbionts, revealing species-species differences which question the current model of ‘one species represents all’, when it comes to mucormycosis.

## Materials and Methods

### Fungal Culture

*R. delemar RA 99-880* and *R. microsporus FP469* were cultured with sabouraud dextrose agar (SDA) or broth (SAB) (10 g/liter mycological peptone, 20 g/liter dextrose), sourced from Sigma-Aldrich, at room temperature. Spores were harvested, 10 days after plating, with phosphate-buffered saline (PBS), centrifuged for 3 min at 3,000 rpm, and washed. Appropriate concentrations of spores were used for further experiments as indicated. To cure spores of their respective bacterial endosymbionts, spores treated with ciprofloxacin were germinated, passaged through sporulation twice, and frozen down for stocks before use (Itabangi *et al*. 2019). Successful curing was confirmed by staining and live cell imaging, as described by Itabangi *et al*. Once cured, spores were subcultured at least twice in ciprofloxacin-free media before use. Strains used in this study are identical to those used by Itabangi *et al*. (Itabangi *et al*. 2019) and Ibrahim *et al*. (Ibrahim *et al*. 2008).

### Macrophage Culture

Macrophages from the J774.A1 cell line were cultured in Dulbecco’s Modified Eagle Medium, (complemented with 10% foetal bovine serum, 1% penicillin, 1% streptomycin and 1% L-glutamine). Macrophages were routinely grown at 37°C, in 5% CO_2_.

### Phagocytosis Assay

Macrophages were incubated for one hour in serum-free DMEM (sfDMEM) prior to infection. Spores were pre-swollen in SAB (2 hr for *R. delemar,* 4 hr for *R. microsporus).* Washed spores were incubated at a 5:1 multiplicity of infection (MOI) with 1 x 10^5^ macrophages as described in Itabangi *et al*. to ensure that >95% of macrophages contained one spore or more. After a 1 hour incubation, excess spores were washed off the surface and the macrophages were incubated for a further 2 hours, before processing for RNA-Seq experiments. For live cell imaging experiments, images were acquired starting immediately after the excess spores were removed.

### Live Cell Imaging

Time course images were taken to determine how LPS pre-treatment (100ng/ml) of macrophages, and Nikkomycin Z pre-treatment (24ug/ml over the course of swelling in SAB) of spores effected phagocytic outcome (Mosser and Zhang 2008). Images were acquired at 20x on a Zeiss Axio Observer, with imaging every 5 minutes. Bright-field and fluorescent images were then analysed for physical lysis using ImageJ V1. Macrophage survival was determined via analysis of the resulting images.

### Genome Sequencing

To confirm the identity of *R. microsporus* (FP469-12.6652333, isolated at the Queen Elizabeth Hospital Birmingham Trauma Centre), DNA was extracted from germinated *R. microsporus* spores following to DNeasy guidelines (Qiagen). Library preparation followed Nextera protocols, and MicrobesNG performed sequencing, utilising an illumina sequencing platform to achieve 30X coverage. The resulting reads were aligned to the indexed genome of *Rhizopus microsporus* (ATCC 52813, accession: GCA_002708625.1).

### Comparative Genomics and Enrichment Analysis

Fishers exact test was used to detect enrichment of Pfam terms between *R. delemar* and *R. microsporus,* terms with a corrected P value of < 0.01 were considered significant. Orthologue genes of *R. delemar* and *R. microsporus* were identified using blast+. R (version 3.3.3) was used to carry out hypergeometric testing of KEGG and GO terms to determine enrichment.

### Bacterial RNA-Seq

Transcriptional analysis of *R. pickettii* was assessed following growth for 4 h at 30 °C, 80-150 rpm in DMEM, or DMEM + cured *R. microsporus* hyphae. Triplicate biological replicates were prepared for each condition. RNA was extracted using the modified Qiagen RNA extraction method. Briefly, TRIzol was used to lyse the samples, which were then either immediately frozen at −20°C and stored for RNA extraction or placed on ice for RNA extraction. After lysis, 0.2 ml of chloroform was added for every 1 ml of TRIzol. Samples were incubated for 3 min, then spun at 12,000 g at 4°C for 15 min. To the aqueous phase, an equal volume of 100% ethanol (EtOH) was added, before the samples were loaded onto RNeasy RNA extraction columns (Qiagen). The manufacturer’s instructions were followed from this point onwards. RNA quality was checked by Agilent, with all RNA integrity number (RIN) scores above 7. One microgram of total RNA was used for cDNA library preparation. Library preparation was done in accordance with the NEBNext pipeline, with libraries quality checked by Agilent. Samples were sequenced using the Illumina NextSeq platform; 150-bp paired-end sequencing was employed (2 x 150 bp) (>10 million reads per sample). The output was aligned to the *R. pickettii* 12J reference with Hisat2 (Version 2.0.5), and downstream analysis was performed with HTSeq (Version 0.10.0) and edgeR (Version 3.16.5).

### Fungal RNA-Seq

RNA was extracted from spores which had either been incubated with the macrophages, or incubated in DMEM for the equivalent time period (3 biological replicates per condition). To remove the macrophages, triton at 1% was used to lyse the macrophages, the resulting solution was then centrifuged for 3 min at 3,000 rpm, and washed, leaving only spores. The DMEM control also received the same treatment. To extract total RNA, the washed samples were immediately immersed in TRIzol and lysed via bead beating at 6,500rpm for 60s. Samples were then either immediately frozen at −20°C and stored for RNA extraction or placed on ice for RNA extraction. After lysis, 0.2 ml of chloroform was added for every 1 ml of TRIzol used in the sample preparation. Samples were incubated for 3 min and then spun at 12,000 *g* at 4°C for 15 min. To the aqueous phase, an equal volume of 100% ethanol (EtOH) was added, before the samples were loaded onto RNeasy RNA extraction columns (Qiagen). The manufacturer’s instructions were followed from this point onwards. RNA quality was checked by Agilent, with all RNA integrity number (RIN) scores above 7 (Schroeder *et al*. 2006). One microgram of total RNA was used for cDNA library preparation. Library preparation was done in accordance with the NEBNext pipeline, with library quality checked by Agilent. Samples were sequenced using the Illumina HiSeq platform; 100-bp paired-end sequencing was employed (2 x 100 bp) (>10 million reads per sample).

### Single Cell RNA-Seq

For single cell sequencing experiments, macrophages were infected with fungal spores, as outlined above. Uninfected macrophages, used as a negative control, were treated in the same manner and underwent mock washes and media changes at identical time points to infected macrophages. Macrophages were isolated and released from the bottom of their wells with accutase, as per manufacturer’s instructions (Technologies n.d.). Once in solution, the macrophages were loaded onto the 10X genomics single cell RNA sequencing pipeline for single cell isolation and library preparation. In total, 1082 single cells were sequenced. The libraries were sequenced on the Illumina Sequencing Platform.

### Data Analysis

For the fungal and bacterial RNA-Seq data, FastQC (version 0.11.5) was employed to ensure the quality of all samples. Hisat2 (version 2.0.5) was used to align reads to the indexed genome of *Rhizopus delemar* RA 99-880 and the indexed genome of *Rhizopus microsporus* (ATCC 52813, accession: GCA_002708625.1) (Ma *et al*. 2009; Mondo *et al*. 2017). HTSeq (version 0.8.0) was used to quantify the output (Anders *et al*. 2015). Trinity and edgeR (version 3.16.5) were then used to analyse differential expression (Robinson *et al*. 2010; Grabherr *et al*. 2011). For the single cell RNA-Seq data, the 10X genomics analysis pipeline (Loupe Cell Browser V 2.0.0, Cell Ranger Version V 2.0.0) was used to align reads to the *mus musculus* genome (version MM10), and quantify the output. For single cell analysis, the samples were then aggregated using the 10X genomics analysis pipeline, to allow comparisons between samples.

## Data Availability

Data have been uploaded to the SRA database and can be accessed via the following accession numbers: PRJNA680574 (*R. microsporus*), PRJNA680572 (*R. delemar*), PRJNA680577 (*R. pickettii*), PRJNA680578 (macrophage).

## Results

### Experimental Design

We set out to investigate paired transcriptional responses of host and fungal cells, whilst also exploring the influence of the endosymbiont on this interaction (Figure 1a). Fungal spores from *Rhizopus delemar* and *Rhizopus microsporus* were either cured via ciprofloxacin treatment to remove the bacterial endosymbiont (cured) or maintained in media permissive to bacterial endosymbiosis (wt). Cured spores were passaged twice in the absence of ciprofloxacin to limit the impact of the drug on transcriptional responses. The cured and wt spores of both *R. delemar* and *R. microsporus* were allowed to swell in sabouraud broth (SAB) until 95% of the population had reached mid isotropic phase. Due to the differences in germination rates between the species (Figure 1b), this occurred at 2 hours for *R. delemar* and 4 hours for *R. microsporus.* Swollen spores were then used to infect the J774.1 murine macrophage-like cell line. Fungal spores were co-cultured with macrophages for one hour, after which unengulfed spores were removed, and phagocytosed spores were incubated within the macrophages for a further two hours. The cells from the resulting infection were processed to explore their transcriptional response to this infection scenario (Figure 1a). Macrophages that had phagocytosed fungal spores were isolated and sequenced via the 10X Genomics Chromium Single Cell Sequencing platform. Macrophages left unexposed to the fungi were used as a negative control. RNA was also isolated from fungal spores (cured and wt) which had been engulfed by macrophages, and this was sequenced with a bulk RNA-Seq approach. Unexposed fungi (cured and wt) were incubated in macrophage media for a matched time and used as a negative control. The resulting data shows the fungal response to phagocytosis by macrophages, as well as the fungal response to the presence of its endosymbiont (Figures 2–5). The macrophage response to the two species is also revealed, a response which appears to differ when the endosymbiont is present for both fungal species (Figures 6–7).

**Figure 1.**
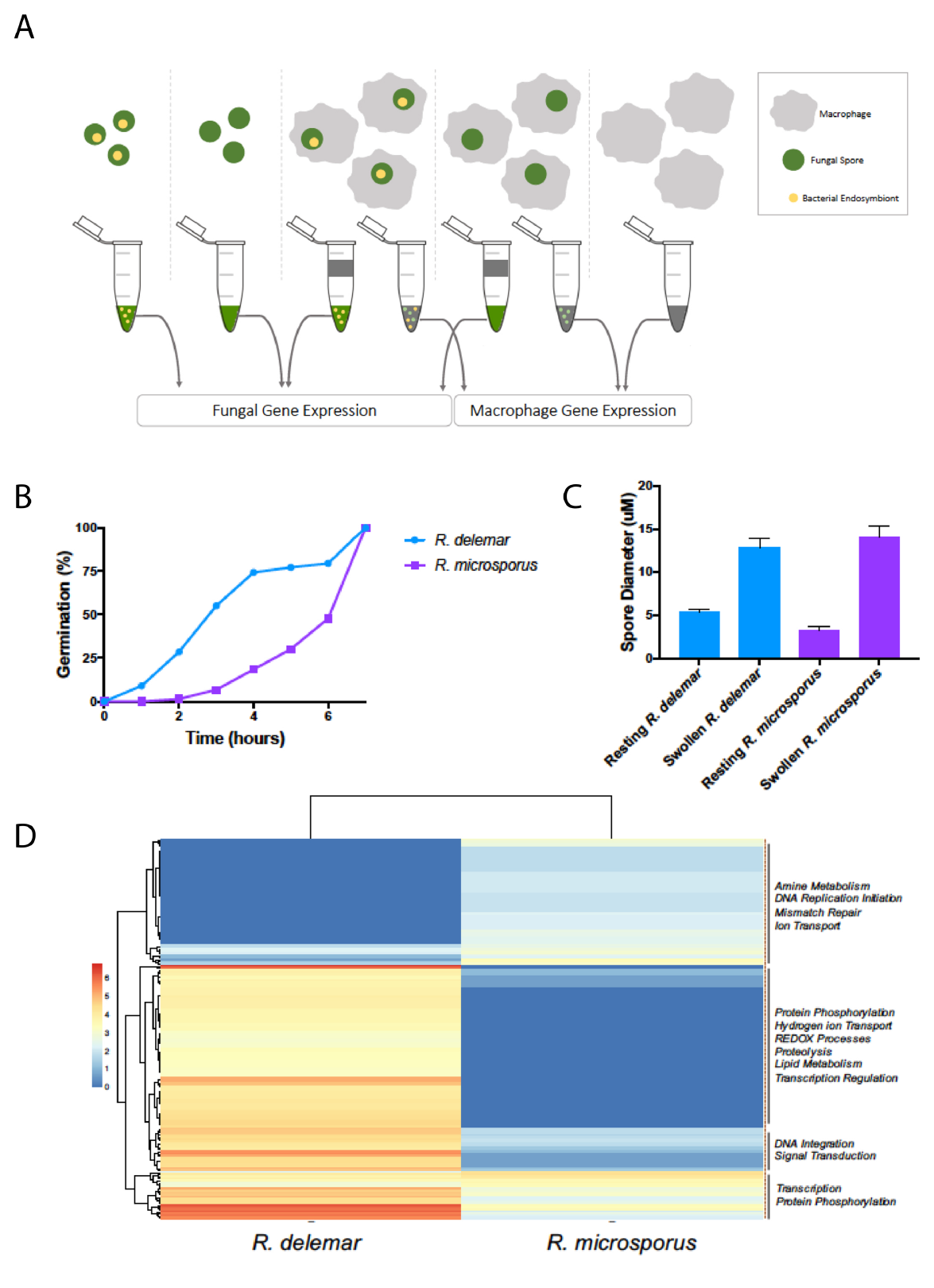
Phenotypic and genomic comparisons. A) Experimental setup: Pre-swollen (SAB) wild-type (wt) or cured fungal spores from *Rhizopus delemar* (2hr) and *Rhizopus microsporus* (4hr) in mid-isotropic phase were co-cultured with J774.1 murine macrophages (sfDMEM) at 5:1 MOI for 1hr, washed to remove unengulfed spores, then co-incubated for a further 2 hours before fungi and host cells were harvested. Pre-swollen spores (SAB) incubated in sfDMEM for 3hr served as the negative control. B) Germination percentage over time for *R. delemar* and *R. microsporus* grown in SAB. C) Spore sizes for *R. delemar* and *R. microsporus* grown in SAB. D) Pfam terms enriched (corrected P < 0.01) in *R. delemar* vs *R. microsporus* genomes, with colour indicating the extent of enrichment (Log_10_ Count).

### Comparative genomics of Rhizopus species

In order to better understand differential disease progression in mucormycosis, we chose to compare *R. delemar* and *R. microsporus*, as they cause a large proportion of mucormycosis infections but appear morphologically dissimilar. *R. delemar* germinates more quickly than *R. microsporus,* with 50% of spores germinating by 3 hours, and a mean spore body size which reaches 12.8 μm over the course of germination (Figure 1b, c). *R. microsporus* germinates at a significantly slower rate, taking 6 hours for 50% of spores to germinate, with a final mean spore body size of 13.9 μm (Figure 1b, c). To better understand their relationship to one another, we compared the gene content of the two species to explore the similarity and differences between them. Previous work has established that the *R. delemar* genome (45.3 Mb, 17,513 genes) is larger, and contains an increased number of genes, compared to the *R. microsporus* genome (26 Mb, 10,959 genes) (Ma *et al*. 2009; Mondo *et al*. 2017). Our results show that, compared to *R. microsporus,* the genome of *R. delemar* is enriched for genes with protein domains (PFAM) associated with ion binding, carbohydrate derivative binding, nucleic acid binding, cytoskeletal protein binding, poly(A) binding, NAD+ ADP-ribosyltransferase activity, protein kinase C activity, translation initiation factor binding and inorganic phosphate transmembrane transporter activity (Supplemental Figure 1, Figure 1d). *R. microsporus* is enriched for genes with protein domains corresponding to nucleoside phosphate binding, early endosome activity and DNA repair complex activity (Supplemental Figure 1, Figure 1d). The clear differences illustrated by genome size and gene content indicate the likelihood of alternative transcriptional responses.

### Alternative transcriptional profiles are presented by R. delemar and R. microsporus in response to innate immune cells

First, we examined overall trends in fungal responses to phagocytosis, obtained through our bulk RNA-Seq approach. We analysed the signal obtained from *R. delemar* and *R. microsporus* samples with principle component analysis (PCA) (Figure 2). We observed large differences between the transcriptomes of both fungal species, when exposed or unexposed to macrophages, while the presence or absence of their respective endosymbionts had a weak but differential effect on PCA. The presence or absence of the endosymbiont appears to have very little bearing on the transcriptional patterns displayed by *R. delemar,* as samples fell into two distinct clusters, most strongly influenced by macrophage exposure (Figure 2a). *R. microsporus* exhibits a similar trend upon exposure to macrophages, however the presence of the endosymbiont also influenced clustering (Figure 2b). Here, we focus on the transcriptional analysis of the host-pathogen-endosymbiont interaction across the two species.

**Figure 2.**
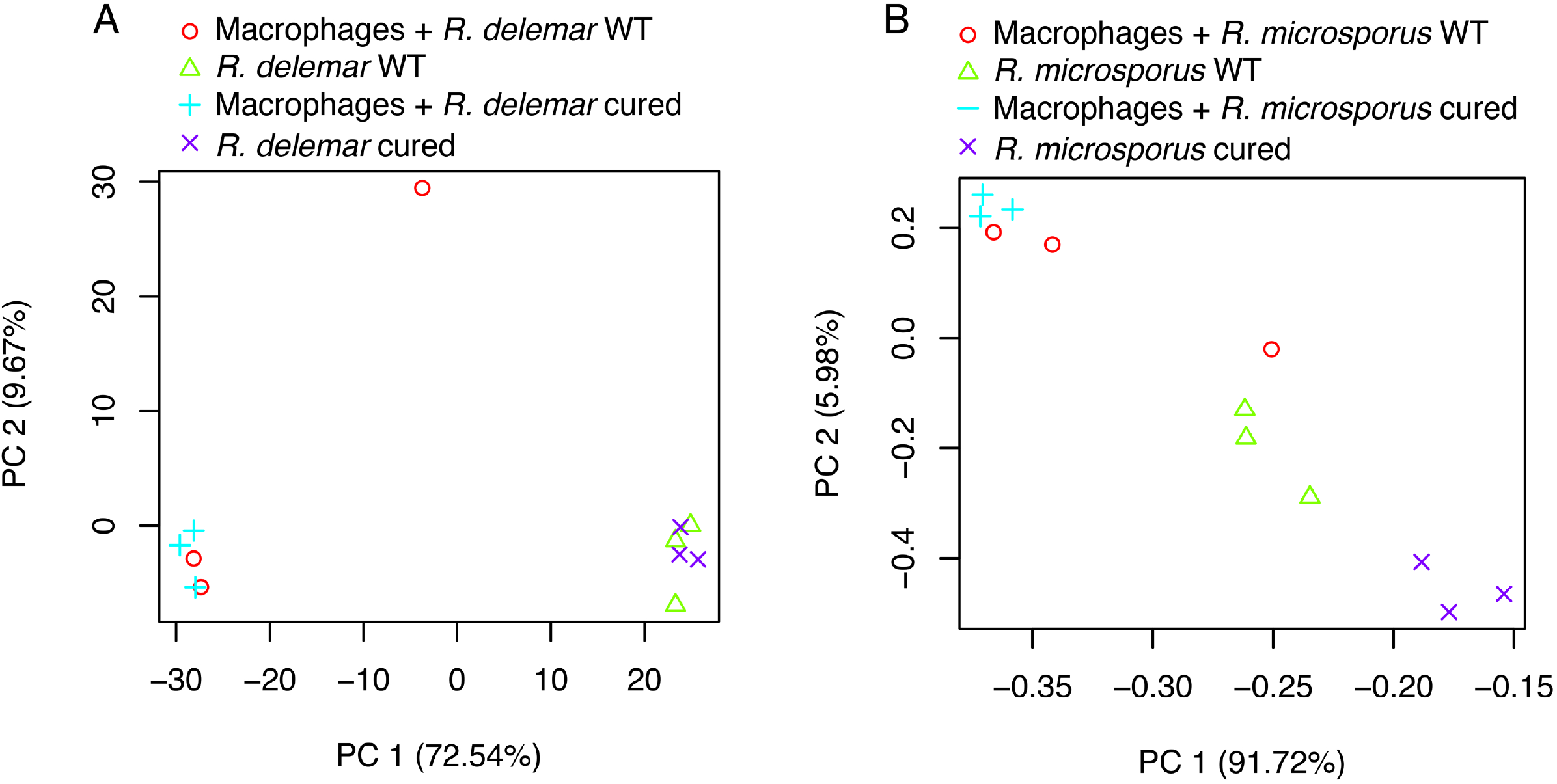
Principal component analysis of fungal genes differentially expressed across all samples. A) *R. delemar* wt and cured, macrophage engulfed or sfDMEM control B) *R. microsporus* wt and cured, macrophage engulfed or sfDMEM control. Biological replicates (n=3) are shown for each sample.

There are 2,493 genes that are significantly differentially expressed (Log fold change > 2; false discovery rate < 0.05) in *R. delemar* across all conditions (Figure 3a), while *R. microsporus* only exhibits 40 genes significantly differentially expressed across all conditions (Log fold change > 2; false discovery rate < 0.05) (Figure 3b). The theme of a muted transcriptional response from *R. microsporus* is also seen within pairwise comparisons of conditions. Pairwise comparisons of differential expression across each experimental condition show similar trends in responses between *R. delemar* and *R. microsporus,* however *R. microsporus* responds with a reduced gene set (Figure 4). Pairwise comparisons showed the biggest shift in transcriptional response is observed when comparing phagocytosed fungal spores to those unexposed to macrophages, regardless of endosymbiont status. When phagocytosed (Supplemental Figure 2), *R. microsporus* upregulates genes enriched in GO categories corresponding to thiamine metabolism, sulfur metabolism, glycerol metabolism, alcohol dehydrogenase activity and transmembrane transporter activity (hypergeometric test, corrected *P* value < 0.05). This is consistent with the fungal response seen to macrophage stress, and the micronutrient scavenging response to nutritional immunity (Parente-Rocha *et al*. 2015; Ballou and Wilson 2016; Andrianaki *et al*. 2018; Shen *et al*. 2018). Phagocytosed *R. microsporus* downregulated genes enriched in GO categories corresponding to rRNA processing, ribosome biogenesis and ribosome localization (hypergeometric test, corrected *P* value < 0.05) (Supplemental Figure 2), consistent with growth arrest within the phagolysosome (Andrianaki *et al*. 2018; Inglesfield *et al*. 2018).

**Figure 3.**
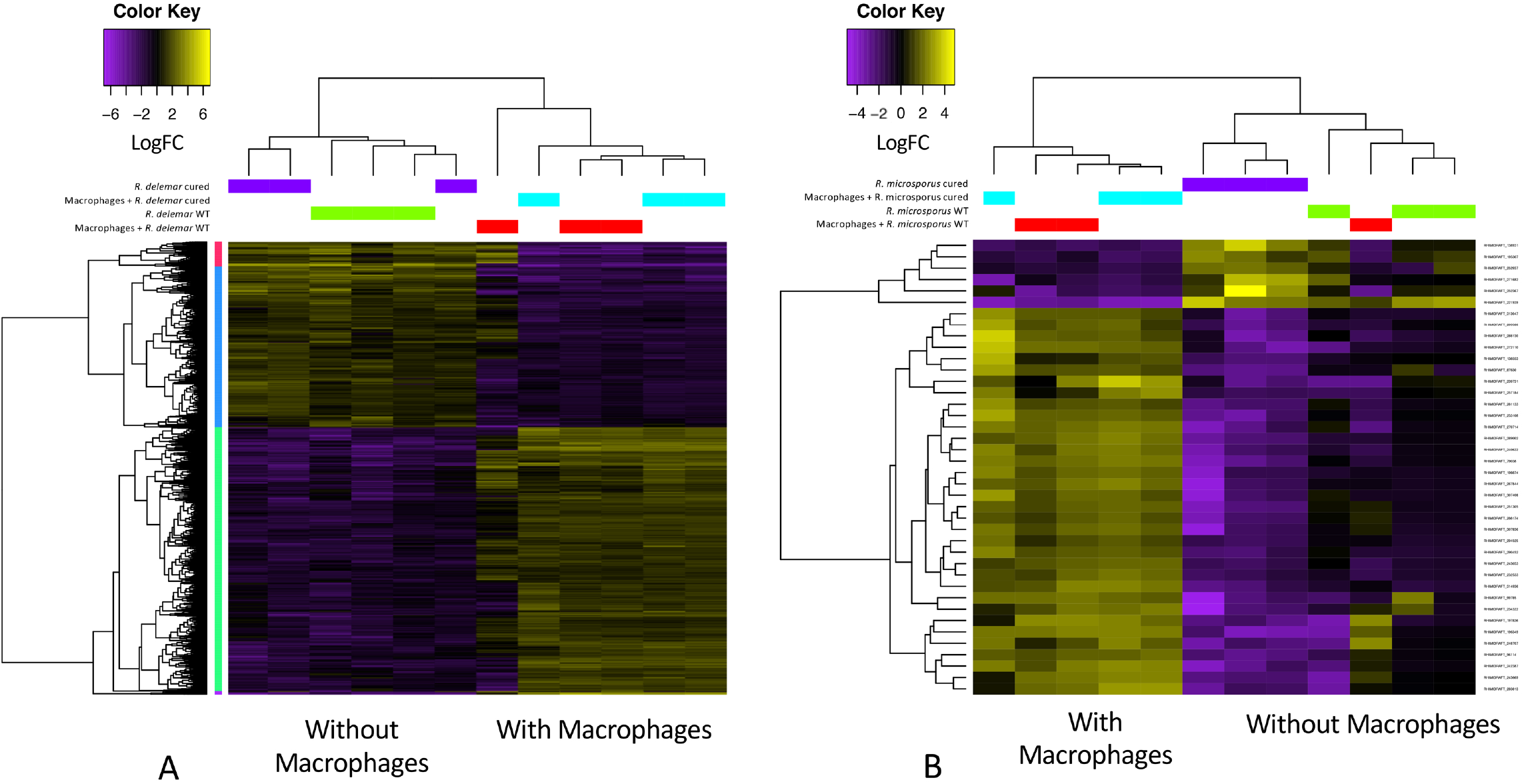
Clustering of fungal transcriptional changes. A) Heatmap displaying differentially expressed genes in *R. delemar.* Expression levels are plotted in Log2, space and meancentered (FDR < 0.001) B) Heatmap displaying differentially expressed genes in *R. microsporus*. Expression levels are plotted in Log2, space and mean-centered (FDR < 0.001).

**Figure 4.**
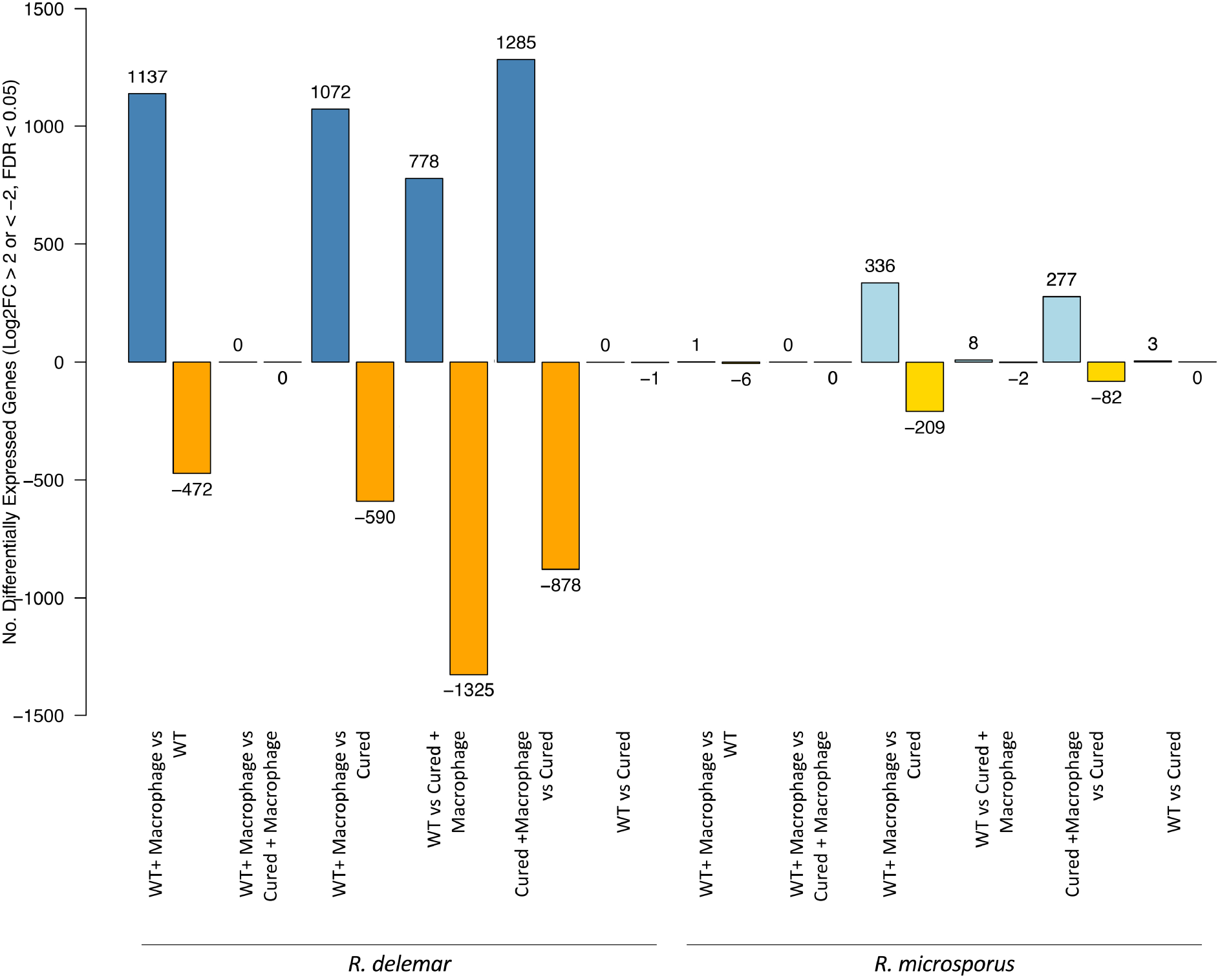
Differential expression of fungal genes in compared conditions. A) The number of genes significantly differentially expressed (multiple corrected P value < 0.05) between samples. Blue bars indicate genes with an increase in expression (LogFC > 2), whilst orange bars indicate genes with a decrease in expression (LogFC < −2).

Comparisons of *R. delemar* conditions reveal that, upon phagocytosis, spores upregulate genes enriched in KEGG classifications corresponding to MAPK signalling, phenylalanine metabolism, tyrosine metabolism, glutathione metabolism and fatty acid synthesis (hypergeometric test, corrected *P* value < 0.05). Upregulation of these processes is consistent with melanin biosynthesis and intra-phagosomal survival (Lorenz and Fink 2001; Yadav *et al*. 2011; Eisenman and Casadevall 2012; Andrianaki *et al*. 2018). Unexposed *R. delemar* spores upregulate genes enriched in KEGG classifications corresponding to ketone body synthesis, protein processing via the endoplasmic reticulum, amino sugar and nucleotide sugar metabolism (hypergeometric test, corrected *P* value < 0.05). This is consistent with metabolic activation and cell wall biogenesis (Figure 5).

**Figure 5.**
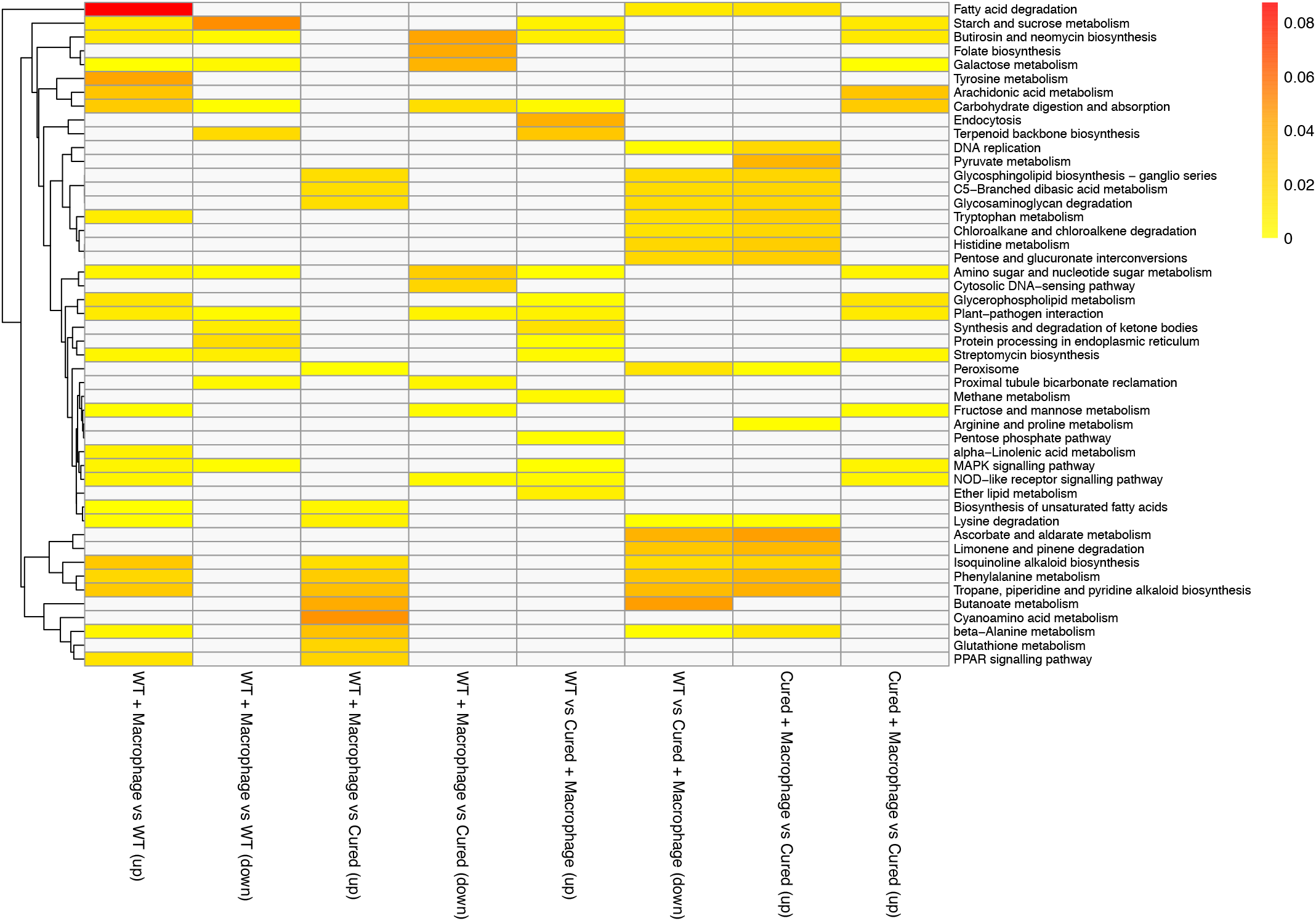
Functions of genes differentially expressed in *R. delemar.* Enriched KEGG categories for the up/down (LogFC > 2 or < −2) regulated genes over sample comparisons. The enrichment of the category is indicated by the colour bar. White corresponds to no enrichment, and yellow to red corresponds to the given P value of the enrichment.

### R. delemar and R. microsporus transcriptional responses to innate immune cells are impacted by loss of endosymbionts

*R. delemar* and *R. microsporus* harbour distinct bacterial endosymbionts (Itabangi *et al*. 2019; Ibrahim *et al*. 2008). The endosymbiont of *R. microsporus* was isolated and identified as *Ralstonia pickettii,* and presence of an unculturable *R. delemar* endosymbiont previously reported by Ibrahim *et al*. was confirmed with PCR (Itabangi *et al*. 2019; Ibrahim *et al*. 2008, data not shown). When comparing transcriptional profiles of wt and cured spores incubated in serum-free DMEM (sfDMEM) for 3 hours (time matched to phagocytosis assay), we observed very few transcriptional changes in either fungal species in response to curing (Supplemental Table 1). In *R. delemar,* a single gene, predicted to be a putative protein phosphatase, was repressed. In *R. microsporus,* three genes were induced: an autophagy-related protein, a C2H2 zinc finger transcription factor, and ribosomal protein L2. In contrast, loss of the endosymbiont significantly impacted the transcriptional responses of both fungal species to macrophages (Figure 4). Specifically, we observed an overall increase in the number of fungal genes differentially regulated upon phagocytosis for both species.

Exposure of wt *R. microsporus* to macrophages induced the expression of one gene with no known function and repressed 6 genes. The repressed genes include an autophagy-related protein and a zinc finger transcription factor, as well as 4 un-annotated genes (Figure 4, Supplemental Table 1). In contrast, when cured *R. microsporus* spores were exposed to macrophages, 277 genes were significantly induced and 82 repressed, compared to unexposed cured spores (Figure 4). Induced genes were enriched (hypergeometric test, corrected *P* value < 0.05) for the following GO categories: organelle organisation, preribosome and ribosome activity, ATPase activity, hydrolase activity, pyrophosphatase activity, helicase activity, nucleic acid binding, RNA metabolism, nitrogen metabolism, chromatin silencing. Repressed genes were enriched (hypergeometric test, corrected *P* value < 0.05) for the following GO categories: oxidoreductase activity, hydrogen sulphide metabolism, glycolysis, sulphur metabolism, hexose catabolism, siderophore activity, iron assimilation, nitrogen metabolism, carboxylic acid metabolism. This suggests an overall failure to properly respond to host stresses such as iron starvation in the absence of the bacterial endosymbiont. Itabangi *et al*. have shown that the presence of the endosymbiont *Ralstonia pickettii* impacts resistance to host-relevant stress and pathogenesis, and this is reflected in the transcriptional changes observed here (Itabangi *et al*. 2019).

A similar impact of the unidentified endosymbiont was observed for *R. delemar.* In wt samples, the overall fungal response to phagocytosis is characterized by a robust transcriptional response: induced genes in wt samples (1137 genes, Figure 4) were enriched for KEGG classifications corresponding to: alanine metabolism, PPAR signalling, aromatic compound biosynthesis and degradation, lysine metabolism, lipid metabolism, MAPK signalling, sugar metabolism, tyrosine metabolism, secondary metabolite biosynthesis (Figure 5). Repressed genes in wt samples (472 genes, Figure 4) were enriched for KEGG classifications corresponding to: carbohydrate metabolism, secondary metabolite biosynthesis, ketone body processing, protein processes, MAPK signalling (Figure 5). In contrast, induced genes in cured samples (1285 genes, Figure 4) were enriched for KEGG classifications corresponding to: Fatty acid metabolism, DNA replication, amino acid metabolism, glycan metabolism, pyruvate metabolism, secondary metabolite processing (Figure 5). Repressed genes in cured samples (878 genes, Figure 4) were enriched for KEGG classifications corresponding to: sugar metabolism, amino acid metabolism, lipid metabolism, MAPK signalling, NOD-like receptor signalling. Again, this suggests that the endosymbiont has an overall suppressive impact on fungal transcription in response to macrophage challenge.

### R. delemar and R. microsporus exhibit distinct transcriptional profiles with implications for virulence

Next, we directly compared the transcriptional patterns of genes shared across the two fungal species. When comparing the transcriptional responses of orthologous genes shared by *R. delemar* and *R. microsporus,* we saw only a small proportion behave similarly (213 genes). When phagocytosed, wt spores from both species upregulate orthologues genes involved in fatty acid catabolism, transcription, regulation via polymerase II, and organelle organization. Phagocytosed cured spores from both species upregulate orthologous genes involved in RNA processing, chromosome organization and condensed chromosome pathways. When unexposed, we see wt spores upregulate orthologous genes involved in translocation, protein binding, siderophore activity, cobalmin processing, and post-translational protein targeting. Cured unexposed spores upregulate orthologous genes with roles in siderophore activity and transferase activity. Overall, *R. delemar* and *R. microsporus* both respond transcriptionally to the presence of macrophages, however the size and composition of this response differs between species. This may reflect the accommodation of different endosymbiont species, as well as the differences in genome size and structure between *R. delemar* and *R. microsporus.*

Finally, we examined the regulation of genes predicted to be involved in ferrous iron transport, as previous work has linked iron scavenging to survival within the phagolysosome (Andrianaki *et al*. 2018). There are 12 genes in the *R. delemar* genome with predicted ferrous iron roles. While 8 showed no significant change over the tested conditions, 3 (R0G3_006623, R0G3_007727, R0G3_011864) appeared highly expressed in wt and cured phagocytosed spores, compared to unexposed spores. The last gene, ROG3_009943, is highly expressed in wt spores unexposed to macrophages. Together, this suggests there may be condition dependent specialization in the expression of ferrous iron transport in *R. delemar.*

### Ralstonia pickettii upregulates fatty acid metabolism and transport in the presence of R. microsporus

To identify the response of *R. microsporus’* bacterial endosymbiont to its fungal host, we analysed transcription of *R. pickettii* isolated from *R. microsporus* in the following conditions: DMEM, and DMEM supplemented with *R. microsporus* substrate (heat killed cured mycelium). When grown in the absence of *R. microsporus* substrate, functions including oxidative phosphorylation, 4Fe-4S binding, heat shock response (HSP20) and cytochrome c activity were upregulated. When supplemented with fungal substrate, an increase in fatty acid metabolism, Acyl-CoA activity, ABC transporter activity and tetR regulation arises, suggestive of fungal substrate use for metabolic processes (Supplemental Figure 3).

### Innate immune cell transcription varies with fungal species and endosymbiont presence

To investigate the innate immune response when challenged with *R. delemar* and *R. microsporus,* we carried out single cell RNA-Seq of J774.A1 murine macrophages, unexposed and exposed for 3 hours to the four types of pre-swollen spores (Figure 1a). Both challenged and unchallenged macrophages separate based on challenge status when DEGs (adjusted P < 0.0001) are used to determine principal components (Supplemental Figure 4a). Enrichment analysis of all DEGs identifies that other than challenge status (Supplemental Figure 4a, PC2) cells strongly differentiate based on cell cycle (Supplemental Figure 4b, PC1). Enrichment analysis of all DEGs identifies the most highly enriched functions to be cell cycle, carbohydrate metabolism, cell signalling, and protein turnover (Supplemental Figure 4b). To identify the transcriptional patterns of genes responding to the spores, we focused on the expression of a subset of genes previously identified as immune response genes (Munoz *et al*. 2018). Principle component analysis of the aggregated transcriptional data supports there is a clear difference in transcription between macrophages that have and have not been exposed to the fungi (Figure 6). Across all exposed conditions, relative to unexposed macrophages, there was a profile consistent with cytokine activation, response to stimulus, and activation of the NF-Kb pathway. This was accompanied by repression of CCL5, which is involved in T-cell recruitment (Figure 7). However, different macrophage profiles can be seen in response to the two fungal species, and these are further influenced by the presence of the endosymbiont. While the response to wt *R. delemar* shows the most deviation from the macrophage-only control, exposure to cured *R. delemar* also elicited a strong and distinct macrophage response (Figures 6,7). Exposure to wt *R. delemar* elicits increased expression of general markers of activation, including GTPase activity and MHC class II protein binding (LAG3 repressor of T-cell activation, H2-M2, IFN-gamma induced IIGP1, MX1, KCTD14, PNP2), growth factor binding, IL1 receptor agonist activity and endocytosis (SERPINE1, ENG, FGFBP3, GM8898, GCNT2, IL1F6) (Martinez *et al*. 2013; Szulzewsky *et al*. 2015). Specifically, we observe modest increases in the expression of IFN-g responsive CXCL10 (3.1 fold) and IRG1/IRG11 (5.9 fold), pro-inflammatory SAA3 (3.5 fold), and ENPP4 (2.8 fold), but also induction of the M2 polarizing PSTPIP2/21 (6.7 fold), the IL-4 responsive signaling modulator CISH (5.4 fold), and the vascular damage responsive F3/F31 (5.9 fold) (Martinez *et al*. 2013; Ye and Sun 2015). These latter genes are not as strongly induced during exposure to *R. microsporus,* which may reflect the aggressive nature of infection by *R. delemar* relative to *R. microsporus.* In contrast, infection with cured *R. delemar* showed a decrease in the induction of these M2-polarisation markers (PSTPIP2, 3.5 fold relative to uninduced). The transcriptional profile is instead shifted to include increased transcription of genes involved in G protein signaling and phosphoinositide binding (PDE7B, CCL1, SCARF1, RGS16, PLEKHA4) (Figure 7).

**Figure 6.**
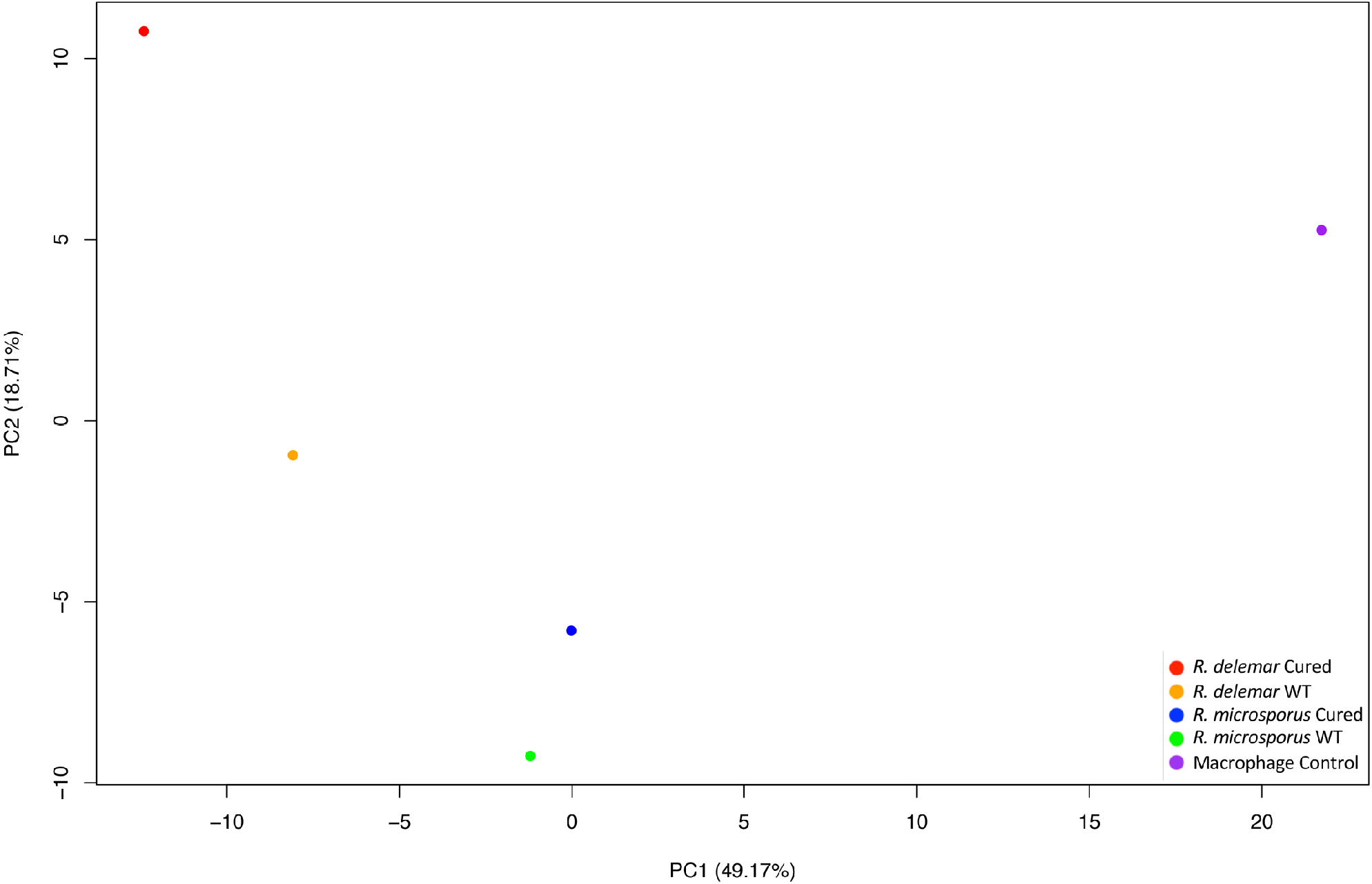
Principal component analysis of macrophage genes differentially expressed across all samples. Single cell sequencing was performed on uninfected and infected macrophages. Transcriptional data from the experiment was analysed with the 10X genomics analysis pipeline, and aggregated prior to principle component analysis.

A similar change in macrophage polarization was observed during exposure to *R. microsporus.* For both wt and cured *R. microsporus,* expression of IFN-g responsive CXCL10, SAA3, and ENPP4 was comparable to unexposed macrophages. Compared to *R. delemar,* exposure to wt *R. microsporus* induced a more limited expression of genes with roles in cytokine activation, ERK1 and ERK2 regulation, and regulation of NF-kB cascade (Figure 7). There was also a weaker induction of the vascular damage responsive F3/F31 genes, and a relatively stronger upregulation of the PLA2G16 phospholipase, TRIM30D, SLC1A2 and the M2 polarizing IL-6 (Berger and Hediger 2006; Luckett-Chastain *et al*. 2017; Raggi *et al*. 2017; van Tol *et al*. 2017; Gerrick *et al*. 2018). However, other key polarizing genes, particularly PSTPIP2/21, SAA3, and ENPP4 were only weakly induced (Supplemental Table 2)(Raggi *et al*. 2017). Overall, this profile suggests a weak M1-like activation consistent with poor phagocytosis and reduced overall antifungal activity that we observe in macrophages interacting with endosymbiontharbouring spores (Itabangi *et al*. 2019).

**Figure 7.**
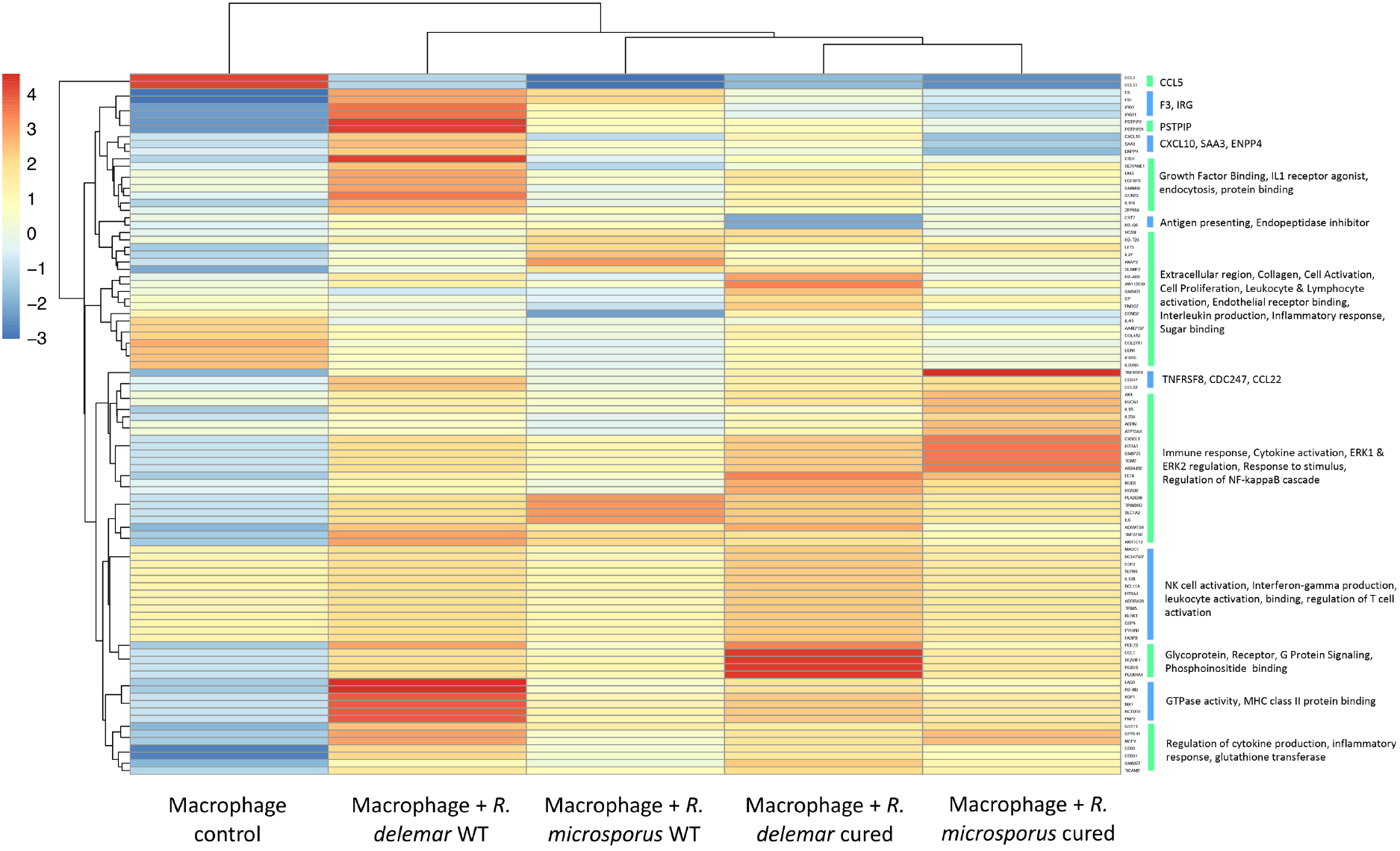
Clustering of macrophage transcription. Heatmap displaying immune response genes significantly differentially expressed between aggregated macrophage populations. Expression levels are plotted in Log2 (FDR < 0.001)

Finally, cured *R. microsporus* induced a strong pro-inflammatory response, which included upregulation of CXCL3, the neutrophil chemoattractant, consistent with our observations of differences in phagocyte recruitment in zebrafish upon infection with wt vs. cured spores (Owen and Mohamadzadeh 2013; Hou *et al*. 2014; Itabangi *et al*. 2019). Markers of NF-kB activation were also strongly induced in this population. Cured *R. microsporus* also strongly induced the expression of TNFRSF8 (CD30), a marker of lymphocyte activation occasionally associated with subcutaneous fungal infections (Muller *et al*. 2011). Overall, this is suggestive of a shift to a more pro-inflammatory profile. Itabangi *et al*. observed that cured *R. microsporus* is more sensitive to phagocyte-mediated killing and phagocyte recruitment compared to wt (Itabangi *et al*. 2019). We therefore went on to test whether a more successful response to the spores could be mounted via the induction of a pro-inflammatory response.

### Chitin synthase inhibition and pro-inflammatory priming regulate infection outcome

*R. delemar* exhibits rapid germination followed by hyphal extension (Sephton-Clark *et al*. 2018). The transcriptional profile we observe here in macrophages exposed to *R. delemar* is consistent with a strong damage response, likely prompted by the germination of *R. delemar* spores. We therefore hypothesised that macrophages may be better able to control the infection if the fungi were slowed in their developmental progress. The necessity of genes involved in chitin synthesis and regulation appeared important for both *R. delemar* and *R. microsporus* in response to phagocytosis (Supplemental Figure 5). In previous work we also highlighted the importance of chitin synthase for germination (Sephton-Clark *et al*. 2018). We therefore tested this model by examining macrophage-fungal interaction following treatment with a chitin synthesis inhibitor. When fungal spores were pre-treated with the chitin synthesis inhibitor Nikkomycin Z (120 μg/ml), both *R. delemar* and *R. microsporus* spores underwent swelling, but failed to germinate, and displayed less chitin/chitosan in their outer cell wall (Supplemental Figure 6). Treatment with lower Nikkomycin Z concentrations (24 μg/ml) allowed spores to swell, but development appears halted after swelling, and macrophage survival increases to 7.5 hours post infection (Figure 8). As the macrophages are better able to control these spores, this suggests that spores undergoing the initial stages of germination may offer less of a challenge for the macrophages than those able to polarise and germinate.

**Figure 8.**
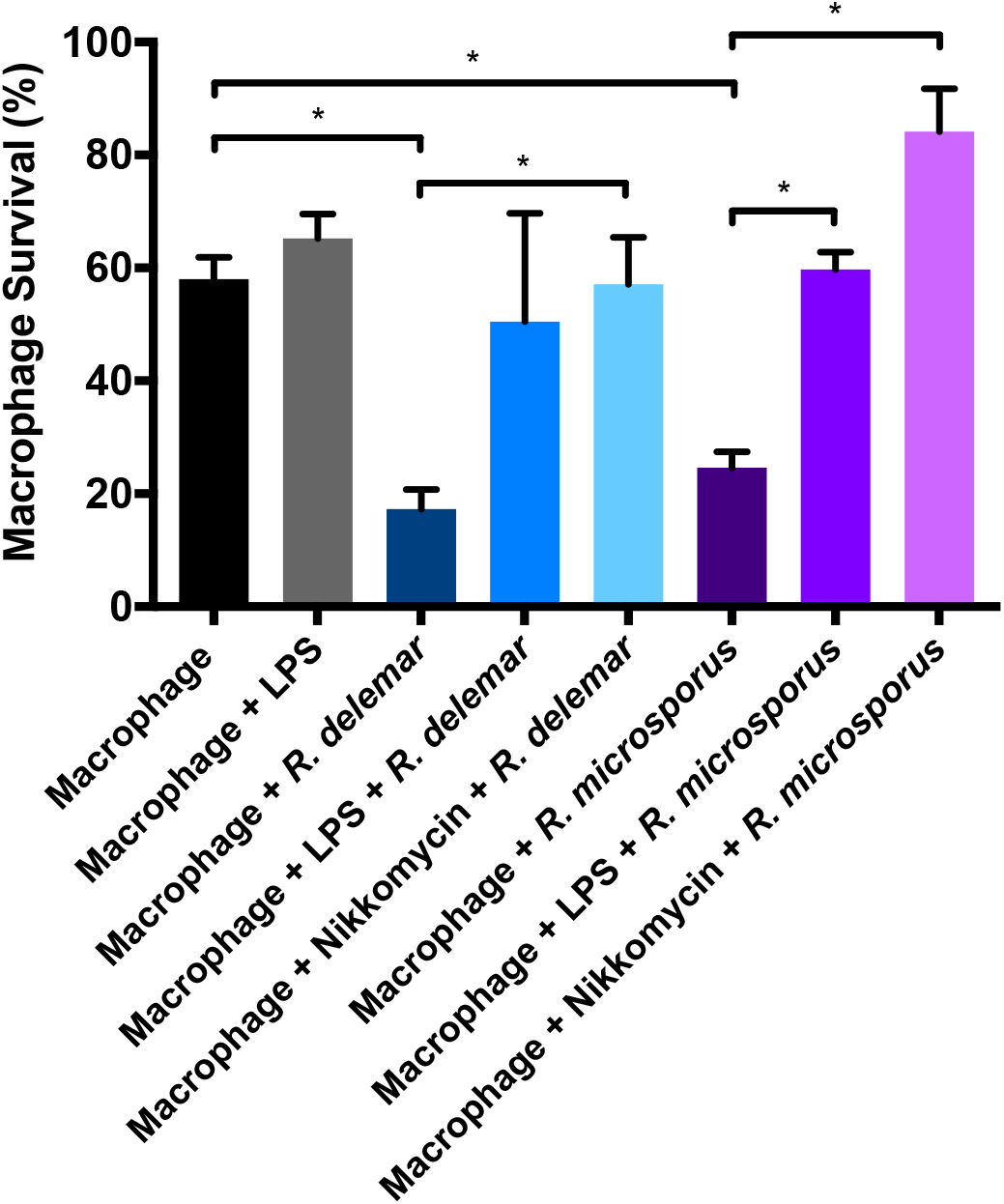
Macrophage survival following exposure to *R. delemar* and *R. microsporus* spores. Macrophages +/− LPS pre-treatment were infected (MOI 5:1) with fungal spores, pre-swollen in SAB consistent with transcriptional experiments. Macrophages were infected with fungal spores that were pre-treated with +/− Nikkomycin Z (24 μg/ml; biological replicate n=3 for each sample). Macrophage survival was determined 7 hr post infection. Significant differences between samples is indicated by (*= p < 0.01). 3 biological replicates per condition.

Our transcriptional data show a strong M2 alternative activation signal during *R. delemar-* macrophage interaction, but a weaker M2 polarisation during *R. microsporus-*macrophage interaction that was further shifted towards NF-kB-mediated M1 upon endosymbiont cure. We therefore hypothesized that shifting the macrophage polarization towards M1 classical activation might have a protective effect upon Mucorales infection. Consistent with this, the pre-treatment of macrophages with NF-kB activating lipopolysaccharide (LPS) significantly improved the ability of macrophages to control *R. microsporus* that maintained the endosymbiont. At 7.5 hours post infection, 59.7% of macrophages survived when pre-treated with LPS, compared to 24.6% without (Figure 8).

## Discussion

This work utilises swollen spores to investigate the interplay between *Rhizopus* species and innate immune cells. Previous studies have used either swollen or resting spores to investigate the interaction between Mucorales species and the host (Inglesfield *et al*. 2018; Andrianaki *et al*. 2018). Itabangi *et al*. demonstrate that swollen *R. microsporus* spores show increased virulence when the endosymbiont *R. pickettii* is present, however, cured swollen and cured resting spores are cleared to a similar extent within *in vivo* models (Itabangi *et al*. 2019). Furthermore, Mucorales spores swell upon immersion in minimal media (Guo *et al*. 1998; Turgeman *et al*. 2013) and likely similarly begin swelling once they are in a host environment. Therefore, understanding the interaction between immune cells and swollen spores is key to understanding pathogenesis. Further transcriptional studies to investigate the differences between interactions of resting and swollen spores with the host would provide useful information on the impact of developmental stage on infection progression.

Whilst the endosymbiont of *R. microsporus* was cultured and identified as *Ralstonia pickettii,* culturing the endosymbiont of *R. delemar* posed a much larger challenge, and this bacterium ultimately remained unculturable. Further efforts to characterise this endosymbiont would offer unique insights into this species-specific symbiosis. Our model uses innate immune cells cultured *in vitro.* Itabangi et al highlight that similar results are seen when using murine macrophages, amoeba, and multiple *in vivo* models (Itabangi *et al*. 2019), and further *in vivo* work will continue to inform our understanding of Mucorales pathogenesis.

In this work we show that the fungal response to innate immune cells differs by species in the Rhizopodaceae family. Although *R. delemar* and *R. microsporus* share a small conserved response on exposure to macrophages, the majority of their response differs. We show that the ability to germinate prior to phagocyte control appears to be key to virulence, as blocking spore germination with the chitin synthase inhibitor Nikkomycin Z improves macrophage survival. A range of germination and virulence phenotypes can be seen throughout the Mucorales, and this highlights a need for further investigation into these differences, to better understand the infections they cause.

We have also shown that the fungal transcriptional response appears largely unperturbed by the presence, or lack, of an endosymbiont, in the absence of stress. However, the presence of an endosymbiont greatly effects the response of and to the host. It has been shown that endosymbionts influence asexual development and sporulation through the regulation of Ras2 (Mondo *et al*. 2017). While we observed limited changes in fungal transcription when comparing wt and cured spores, we did observe changes in the expression of fungal genes upon exposure to macrophages. Itabangi et al show that the endosymbiont of *R. microsporus* enhances virulence through the secretion of an anti-phagocytic factor. The endosymbiont also impacts organisation of the fungal plasma membrane, as well as environmental stress resistance and resistance to macrophage-mediated killing (Itabangi *et al*. 2019). Here, we show that this is mirrored by the transcriptional response of the wt and cured spores of both *Rhizopus* species to macrophages, and in the macrophage response to infection.

As anticipated, activation of pro-inflammatory pathways increased macrophage survival in response to the spores. This consolidates several studies which demonstrate improved innate immune cell response to fungal pathogens when primed (Blasi *et al*. 1995; Municio *et al*. 2013; Rogers *et al*. 2013). It has been shown that the early immune response to a *Mucor circinelloides* infection is dependent on the formation of a pro-inflammatory TNF-α-expressing granuloma-like structure that controls but does not kill spores in the zebrafish model (Inglesfield *et al*. 2018). Ungerminated *Rhizopus* spores are highly resistant to ROS/RNS, which may mediate survival within granulomata.

Interestingly, we see the chitin synthase inhibitor, Nikkomycin Z, is able to inhibit germination in both *R. delemar* and *R. microsporus.* This demonstrates the requirement of chitin synthase for spore development in these species, reflected in our transcriptional analysis. Treated spores remain viable and swell, but do not polarise. The ability to germinate also has consequences for virulence. The ability of macrophages to control swollen spores, but not subsequently polarised ones, highlights that the developmental stage of the spore is key to innate immune success and control. Whilst many fungal pathogens are virulent in an ungerminated form, germination appears a key virulence factor for these filamentous fungi. This is confirmed by work which shows that the developmental stage of *R. microsporus, R. delemar* and *Rhizopus oryzae* spores impacts pathogenesis in zebrafish and murine models (Andrianaki *et al*. 2018; Itabangi *et al*. 2019). Specifically, infection with pre-swollen spores leads to evasion of macrophage-mediated immunity and increased pathogenesis (Andrianaki *et al*. 2018; Itabangi *et al*. 2019). In both models, spore clearance is dependent on phagocyte recruitment. In addition, phagocyte recruitment (Itabangi *et al*. 2019) and activation (Figure 7) is influenced by endosymbiont status (Andrianaki *et al*. 2018; Itabangi *et al*. 2019). Finally, we extended this analysis by revealing profound differences in the host response to two closely related *Rhizopus* species. In particular, we observe a M2/damage-associated response during infection with wt *R. delemar* spores that is shifted towards an M1 protective response upon infection with cured *R. microsporus* spores. We reinforce this finding through experimental modulation of macrophage polarization, showing that exposure to strongly M1-polarizing LPS is sufficient to reduce macrophage killing by wt *R. delemar* spores. Therefore, our data provide a framework for beginning to understand differences in the relative virulence of pathogenic Mucorales species and underpin our finding of cross-kingdom fungal-bacterial symbiosis influencing mammalian disease.

## Author contributions

PSC conceived and designed the experiments, collected the data, performed the analysis and interpretation, and wrote the manuscript.

JFM, HI, KV and CAC contributed to interpretation of the data, and contributed to the manuscript.

ERB conceived and designed the experiments, contributed to interpretation of the data, and wrote the manuscript.

## Funding

PSC was supported by a BBSRC MIBTP PhD Studentship (BB/M01116X/1). This work was supported by a Wellcome Trust Seed award to KV (108387/Z/15/Z). HI was supported by the Wellcome Trust Strategic Award in Medical Mycology and Fungal Immunology (097377). CAC and JFM were funded by the National Institute of Allergy and Infectious Diseases, National Institutes of Health, under award U19AI110818 to the Broad Institute. ERB was supported by the UK Biotechnology and Biological Research Council (BB/M014525/1) and a Sir Henry Dale Fellowship jointly funded by the Wellcome Trust and the Royal Society (211241/Z/18/Z).

## Acknowledgments

We are grateful to the University of Birmingham’s Genomics Services Facility and to Deborah Croom-Carter for her technical support.

---

**Supplemental Figure 1** Table of Pfam terms found to be enriched (corrected P < 5 x 10^-8^) in *R. delemar* and *R. microsporus* upon genome comparison.

**Supplemental Figure 2 Clustering of fungal transcription with GO annotation**. Heatmap displaying all differentially expressed genes, across all conditions, in *R. microsporus.* Expression levels are plotted in Log2, space and mean-centered (FDR < 0.001). GeneIDs correspond to the genome of *Rhizopus microsporus* (ATCC 52813, accession: GCA_002708625.1)

**Supplemental Figure 3 *Ralstonia picketti* expression.** Volcano plot highlighting significantly (red, FDR < 0.0001) differentially expressed genes from *Ralstonia picketti* grown in DMEM vs DMEM supplemented with *R. microsporus* substrate. Labelled genes show a LogFC ± 2 between these two conditions. Gene IDs refer to the *R. pickettii* 12J reference genome.

**Supplemental Figure 4 DEGs and clustering of J774.1 macrophages**. A) Principal component analysis of single cells based on significantly differentially expressed genes (top), and immune response genes (bottom). B) Gene counts per million (Log10(CPM+1)) for all genes differentially expressed (adjusted p value < 0.0001), per cell, with genes falling into enriched functions labelled by colour. 1082 single cells spanning all conditions represented.

**Supplemental Figure 5 Expression of chitin synthase genes**. A) Heatmap showing the expression of chitin synthase genes in *R. delemar.* B) Heatmap showing the expression of chitin synthase genes in *R. microsporus.*

**Supplemental Figure 6 Chitin synthase inhibition of *R. delemar* and *R. microsporus* germination**. *R. delemar* and *R. microsporus* treated with Nikkomycin Z in SAB at indicated concentrations. Fluorescence indicates calcofluor white staining, and thus the availability of chitin/chitosan in the cell wall.

**Supplemental Table 1** Selected genes differentially regulated in pairwise comparisons, with Pfam annotations provided where available.

**Supplemental Table 2** Table displaying the LogFC values for a subset of genes displayed in Figure 7.

**Supplemental Table 3** Table displaying genes differentially expressed by *R. delemar* (2,493), as displayed in Figure 3a.

## Notes

### Competing Interest Statement

The authors have declared no competing interest.

### Summary of Updates

The text has been revised to reflect reviewer feedback and an additional dataset for Ralstonia pickettii has been added. Accession numbers for all datasets are now updated.

## References

Alan, E. C. (2019). “An Emergent Entity: Indolent Mucormycosis of the Paranasal Sinuses, A Multicenter Study.” 2193: 92–100.

Anders, S., P. T. Pyl and W. Huber (2015). “HTSeq--a Python framework to work with high-throughput sequencing data.” Bioinformatics 31(2): 166–169.

Andrianaki, A.M., I. Kyrmizi, K. Thanopoulou, C. Baldin, E. Drakos, S. S. M. Soliman, A. C. Shetty, C. McCracken, T. Akoumianaki, K. Stylianou, P. Ioannou, C. Pontikoglou, H. A. Papadaki, M. Tzardi, V. Belle, E. Etienne, A. Beauvais, G. Samonis, D. P. Kontoyiannis, E. Andreakos, V. M. Bruno, A. S. Ibrahim and G. Chamilos (2018). “Author Correction: Iron restriction inside macrophages regulates pulmonary host defense against Rhizopus species.” Nat Commun 9(1): 5015.

Baldin, C. and A. S. Ibrahim (2017). “Molecular mechanisms of mucormycosis-The bitter and the sweet.” PLoS Pathog 13(8): e1006408.

Ballou, E.R. and D. Wilson (2016). “The roles of zinc and copper sensing in fungal pathogenesis.” Curr Opin Microbiol 32: 128–134.

Berger, U. V. and M. A. Hediger (2006). “Distribution of the glutamate transporters GLT-1 (SLC1A2) and GLAST (SLC1A3) in peripheral organs.” Anat Embryol (Berl) 211(6): 595–606.

Blasi, E., R. Barluzzi, R. Mazzolla, B. Tancini, S. Saleppico, M. Puliti, L. Pitzurra and F. Bistoni (1995). “Role of nitric oxide and melanogenesis in the accomplishment of anticryptococcal activity by the BV-2 microglial cell line.” J Neuroimmunol 58(1): 111–116.

Carter ME, Carpenter SCD, Dubrow ZE, Sabol MR, Rinaldi FC, Lastovetsky OA, Mondo SJ, Pawlowska TE, Bogdanove AJ (2020). “A TAL effector-like protein of an endofungal bacterium increases the stress tolerance and alters the transcriptome of the host.” Proc Natl Acad Sci U S A 117(29):17122–17129.

Chamilos, G., R. E. Lewis, G. Lamaris, T. J. Walsh and D. P. Kontoyiannis (2008). “Zygomycetes hyphae trigger an early, robust proinflammatory response in human polymorphonuclear neutrophils through toll-like receptor 2 induction but display relative resistance to oxidative damage.” Antimicrob Agents Chemother 52(2): 722–724.

Eisenman, H. C. and A. Casadevall (2012). “Synthesis and assembly of fungal melanin.” Appl Microbiol Biotechnol 93(3): 931–940.

Estrada-de los Santos P, Palmer M, Chávez-Ramírez B, Beukes C, Steenkamp ET, Briscoe L, Khan N, Maluk M, Lafos M, Humm E, et al (2018). “Whole genome analyses suggests that Burkholderia sensu lato contains two additional novel genera (Mycetohabitans gen. nov., and Trinickia gen. nov.): Implications for the evolution of diazotrophy and nodulation in the Burkholderiaceae.” Genes (Basel) 9(8):389.

Gerrick, K. Y., E. R. Gerrick, A. Gupta, S. J. Wheelan, S. Yegnasubramanian and E. M. Jaffee (2018). “Transcriptional profiling identifies novel regulators of macrophage polarization.” PLoS One 13(12): e0208602.

Ghuman, H. and K. Voelz (2017). “Innate and Adaptive Immunity to Mucorales.” J Fungi (Basel) 3(3).

Grabherr, M. G., B. J. Haas, M. Yassour, J. Z. Levin, D. A. Thompson, I. Amit, X. Adiconis, L. Fan, R. Raychowdhury, Q. Zeng, Z. Chen, E. Mauceli, N. Hacohen, A. Gnirke, N. Rhind, F. di Palma, B. W. Birren, C. Nusbaum, K. Lindblad-Toh, N. Friedman and A. Regev (2011). “Full-length transcriptome assembly from RNA-Seq data without a reference genome.” Nat Biotechnol 29(7): 644–652.

Hou, Y., H. Lin, L. Zhu, Z. Liu, F. Hu, J. Shi, T. Yang, X. Shi, H. Guo, X. Tan, L. Zhang, Q. Wang, Z. Li and Y. Zhao (2014). “The inhibitory effect of IFN-gamma on protease HTRA1 expression in rheumatoid arthritis.” J Immunol 193(1): 130–138.

Ibrahim, A. S., B. Spellberg and J. Edwards, Jr. (2008). “Iron acquisition: a novel perspective on mucormycosis pathogenesis and treatment.” Curr Opin Infect Dis 21(6): 620–625.

Inglesfield, S., A. Jasiulewicz, M. Hopwood, J. Tyrrell, G. Youlden, M. Mazon-Moya, O. R. Millington, S. Mostowy, S. Jabbari and K. Voelz (2018). “Robust Phagocyte Recruitment Controls the Opportunistic Fungal Pathogen Mucor circinelloides in Innate Granulomas In Vivo.” MBio 9(2).

Itabangi, H., P. Sephton-Clark, X. Zhou, I. Insua, M. Probert, J. Correia, P. Moynihan, T. Gebremariam, Y. Gu, L. Lin, A. S. Ibrahim, G. D. Brown, J. S. King, F. Fernandez Trillo, E. R. Ballou and K. Voelz (2019). “A bacterial endosymbiont enables fungal immune evasion during fatal mucormycete infection “ BioRXiv.

Kraibooj, K., H. R. Park, H. M. Dahse, C. Skerka, K. Voigt and M. T. Figge (2014). “Virulent strain of Lichtheimia corymbifera shows increased phagocytosis by macrophages as revealed by automated microscopy image analysis.” Mycoses 57 Suppl 3: 56–66.

Liu, M., L. Lin, T. Gebremariam, G. Luo, C. D. Skory, S. W. French, T. F. Chou, J. E. Edwards, Jr. and A. S. Ibrahim (2015). “Fob1 and Fob2 Proteins Are Virulence Determinants of Rhizopus oryzae via Facilitating Iron Uptake from Ferrioxamine.” PLoS Pathog 11(5): e1004842.

Lorenz, M. C. and G. R. Fink (2001). “The glyoxylate cycle is required for fungal virulence.” Nature 412(6842): 83–86.

Luckett-Chastain, L. R., M. L. Cottrell, B. M. Kawar, M. A. Ihnat and R. M. Gallucci (2017). “Interleukin (IL)-6 modulates transforming growth factor-beta receptor I and II (TGF-betaRI and II) function in epidermal keratinocytes.” Exp Dermatol 26(8): 697–704.

Ma, L.-J., A. S. Ibrahim, C. Skory, M. G. Grabherr, G. Burger, M. Butler, M. Elias, A. Idnurm, B. F. Lang, T. Sone, A. Abe, S. E. Calvo, L. M. Corrochano, R. Engels, J. Fu, W. Hansberg, J.-M. Kim, C. D. Kodira, M. J. Koehrsen, B. P. Liu, D. Miranda-Saavedra, S. O&apos;leary, L. Ortiz-Castellanos, R. Poulter, J. Rodriguez-Romero, J. Ruiz-Herrera, Y.-Q. Shen, Q. Zeng, J. Galagan, B. W. Birren, C. A. Cuomo and B. L. Wickes (2009). “Genomic Analysis of the Basal Lineage Fungus Rhizopus oryzae Reveals a Whole-Genome Duplication.” PLoS genetics 5(7): e1000549.

Martinez, F. O., L. Helming, R. Milde, A. Varin, B. N. Melgert, C. Draijer, B. Thomas, M. Fabbri, A. Crawshaw, L. P. Ho, N. H. Ten Hacken, V. Cobos Jimenez, N. A. Kootstra, J. Hamann, D. R. Greaves, M. Locati, A. Mantovani and S. Gordon (2013). “Genetic programs expressed in resting and IL-4 alternatively activated mouse and human macrophages: similarities and differences.” Blood 121(9): e57–69.

Mendoza, L., R. Vilela, K. Voelz, A. S. Ibrahim, K. Voigt and S. C. Lee (2014). “Human Fungal Pathogens of Mucorales and Entomophthorales.” Cold Spring Harb Perspect Med 5(4).

Mondo, S. J., O. A. Lastovetsky, M. L. Gaspar, N. H. Schwardt, C. C. Barber, R. Riley, H. Sun, I. V. Grigoriev and T. E. Pawlowska (2017). “Bacterial endosymbionts influence host sexuality and reveal reproductive genes of early divergent fungi.” Nat Commun 8(1): 1843.

Mosser, D. M. and X. Zhang (2008). “Activation of murine macrophages.” Curr Protoc Immunol Chapter 14: Unit 14 12.

Muller, C. S., R. Schmaltz, T. Vogt and C. Pfohler (2011). “Expression of CD30 antigen in superficial mycotic infections of the skin: a possible non-specific finding without clinical relevance.” Mycoses 54(5): e360–363.

Municio, C., Y. Alvarez, O. Montero, E. Hugo, M. Rodriguez, E. Domingo, S. Alonso, N. Fernandez and M. S. Crespo (2013). “The response of human macrophages to beta-glucans depends on the inflammatory milieu.” PLoS One 8(4): e62016.

Munoz, J. F., T. Delorey, C. B. Ford, B. Y. Li, D. A. Thompson, R. P. Rao and C. A. Cuomo (2018). “Coordinated host-pathogen transcriptional dynamics revealed using sorted subpopulations and single, Candida albicans infected macrophages.” BioRXiv.

Owen, J. L. and M. Mohamadzadeh (2013). “Macrophages and chemokines as mediators of angiogenesis.” Front Physiol 4: 159.

Parente-Rocha, J. A., A. F. Parente, L. C. Baeza, S. M. Bonfim, O. Hernandez, J. G. McEwen, A. M. Bailao, C. P. Taborda, C. L. Borges and C. M. Soares (2015). “Macrophage Interaction with Paracoccidioides brasiliensis Yeast Cells Modulates Fungal Metabolism and Generates a Response to Oxidative Stress.” PLoS One 10(9): e0137619.

Raggi, F., S. Pelassa, D. Pierobon, F. Penco, M. Gattorno, F. Novelli, A. Eva, L. Varesio, M. Giovarelli and M. C. Bosco (2017). “Regulation of Human Macrophage M1-M2 Polarization Balance by Hypoxia and the Triggering Receptor Expressed on Myeloid Cells-1.” Front Immunol 8: 1097.

Robinson, M. D., D. J. McCarthy and G. K. Smyth (2010). “edgeR: a Bioconductor package for differential expression analysis of digital gene expression data.” Bioinformatics 26(1): 139–140.

Rogers, H., D. W. Williams, G. J. Feng, M. A. Lewis and X. Q. Wei (2013). “Role of bacterial lipopolysaccharide in enhancing host immune response to Candida albicans.” Clin Dev Immunol 2013: 320168.

Schmidt, S., L. Tramsen, S. Perkhofer, C. Lass-Florl, M. Hanisch, F. Roger, T. Klingebiel, U. Koehl and T. Lehrnbecher (2013). “Rhizopus oryzae hyphae are damaged by human natural killer (NK) cells, but suppress NK cell mediated immunity.” Immunobiology 218(7): 939–944.

Schroeder, A., O. Mueller, S. Stocker, R. Salowsky, M. Leiber, M. Gassmann, S. Lightfoot, W. Menzel, M. Granzow and T. Ragg (2006). “The RIN: an RNA integrity number for assigning integrity values to RNA measurements.” BMC Mol Biol 7: 3.

Sephton-Clark, P. C. S., J. F. Munoz, E. R. Ballou, C. A. Cuomo and K. Voelz (2018). “Pathways of Pathogenicity: Transcriptional Stages of Germination in the Fatal Fungal Pathogen Rhizopus delemar.” mSphere 3(5).

Shen, Q., M. J. Beucler, S. C. Ray and C. A. Rappleye (2018). “Macrophage activation by IFN-gamma triggers restriction of phagosomal copper from intracellular pathogens.” PLoS Pathog 14(11): e1007444.

Spellberg, B., J. Edwards, Jr. and A. Ibrahim (2005). “Novel perspectives on mucormycosis: pathophysiology, presentation, and management.” Clin Microbiol Rev 18(3): 556–569.

Szulzewsky, F., A. Pelz, X. Feng, M. Synowitz, D. Markovic, T. Langmann, I. R. Holtman, X. Wang, B. J. Eggen, H. W. Boddeke, D. Hambardzumyan, S. A. Wolf and H. Kettenmann (2015). “Glioma-associated microglia/macrophages display an expression profile different from M1 and M2 polarization and highly express Gpnmb and Spp1.” PLoS One 10(2): e0116644.

van Tol, S., A. Hage, M. I. Giraldo, P. Bharaj and R. Rajsbaum (2017). “The TRIMendous Role of TRIMs in Virus-Host Interactions.” Vaccines (Basel) 5(3).

Warris, A., M. G. Netea, P. E. Verweij, P. Gaustad, B. J. Kullberg, C. M. Weemaes and T. G. Abrahamsen (2005). “Cytokine responses and regulation of interferon-gamma release by human mononuclear cells to Aspergillus fumigatus and other filamentous fungi.” Med Mycol 43(7): 613–621.

Yadav, A. K., P. R. Desai, M. N. Rai, R. Kaur, K. Ganesan and A. K. Bachhawat (2011). “Glutathione biosynthesis in the yeast pathogens Candida glabrata and Candida albicans: essential in C. glabrata, and essential for virulence in C. albicans.” Microbiology 157(Pt 2): 484–495.

Ye, R. D. and L. Sun (2015). “Emerging functions of serum amyloid A in inflammation.” J Leukoc Biol 98(6): 923–929.

