## Supplemental figures for "Bacterial endosymbionts influence fungal transcriptional profiles with implications for host response in the human fungal pathogens *Rhizopus microsporus* and *Rhizopus delemar*": Supp-1.pdf

| | *R. delemar* | *R. microsporus* | q-value |
|---|---|---|---|
| PF01498 | 873 | 1 | 6.96E-127 |
| PF07714 | 418 | 0 | 4.69E-60 |
| PF00385 | 499 | 8 | 7.87E-60 |
| PF13358 | 532 | 32 | 3.27E-42 |
| PF03372 | 373 | 10 | 1.17E-39 |
| PF08238 | 284 | 2 | 1.47E-36 |
| PF13551 | 258 | 3 | 1.62E-31 |
| PF00400 | 429 | 31 | 1.59E-30 |
| PF00078 | 330 | 14 | 2.18E-30 |
| PF13893 | 293 | 9 | 7.33E-30 |
| PF07282 | 10 | 78 | 1.09E-28 |
| PF00077 | 177 | 0 | 1.58E-24 |
| PF13516 | 194 | 2 | 7.20E-24 |
| PF13606 | 179 | 1 | 4.45E-23 |
| PF12895 | 164 | 0 | 1.15E-22 |
| PF13894 | 374 | 36 | 1.49E-21 |
| PF08284 | 140 | 1 | 2.36E-17 |
| PF07653 | 153 | 3 | 9.95E-17 |
| PF00271 | 124 | 1 | 4.77E-15 |
| PF12894 | 2 | 33 | 1.74E-13 |
| PF13833 | 94 | 0 | 3.81E-12 |
| PF02992 | 126 | 4 | 3.94E-12 |
| PF13191 | 4 | 33 | 8.65E-12 |
| PF09668 | 99 | 1 | 1.51E-11 |
| PF13857 | 99 | 1 | 1.51E-11 |
| PF14259 | 87 | 0 | 3.94E-11 |
| PF00096 | 61 | 85 | 6.25E-11 |
| PF03732 | 129 | 6 | 6.41E-11 |
| PF07690 | 151 | 10 | 7.81E-11 |
| PF13041 | 0 | 23 | 8.70E-11 |
| PF13812 | 92 | 1 | 1.45E-10 |
| PF12937 | 42 | 69 | 1.56E-10 |
| PF13504 | 80 | 0 | 2.17E-10 |
| PF12762 | 95 | 2 | 6.02E-10 |
| PF13637 | 120 | 6 | 7.56E-10 |
| PF00153 | 177 | 17 | 8.42E-10 |
| PF00512 | 76 | 0 | 8.48E-10 |
| PF08662 | 93 | 2 | 8.84E-10 |
| PF04670 | 89 | 2 | 3.54E-09 |
| PF01209 | 82 | 1 | 3.55E-09 |
| PF00025 | 115 | 6 | 3.82E-09 |
| PF13374 | 70 | 0 | 5.60E-09 |
| PF13650 | 86 | 2 | 8.34E-09 |
| PF00665 | 87 | 2 | 8.97E-09 |
| PF05148 | 77 | 1 | 1.28E-08 |
| PF08659 | 66 | 0 | 2.28E-08 |
| PF12773 | 0 | 18 | 2.66E-08 |
| PF03184 | 64 | 0 | 3.66E-08 |
| PF09011 | 65 | 0 | 3.66E-08 |
| PF00071 | 18 | 40 | 4.49E-08 |
