## Supplemental figures for "Bacterial endosymbionts influence fungal transcriptional profiles with implications for host response in the human fungal pathogens *Rhizopus microsporus* and *Rhizopus delemar*": Supp-Table1.pdf

**Table S1: Selected genes differentially regulated in pairwise comparisons**

| | WT vs. Cured | WT + Macrophage vs. WT | Cured + Macrophage vs. Cured | Annotation |
|---|---|---|---|---|
| ***R. delemar*** | | | | |
| RO3G_009519 | down >log2 fold | no change | no change | Protein phosphatase |
| ***R. microsporus*** | | | | |
| RHIMIDRAFT_279423 | Up >log2 fold | down >log2 fold | no change | C2H2 type zinc-finger |
| RHIMIDRAFT_95344 | Up >log2 fold | down >log2 fold | no change | Autophagy-related C terminal domain |
| RHIMIDRAFT_292382 | no change | down >log2 fold | no change | no annotation |
| RHIMIDRAFT_206880 | no change | down >log2 fold | no change | no annotation |
| RHIMIDRAFT_199741 | no change | down >log2 fold | down >log2 fold | no annotation |
| RHIMIDRAFT_84646 | no change | down >log2 fold | no change | no annotation |
| RHIMIDRAFT_197836 | no change | Up >log2 fold | no change | no annotation |
| RHIMIDRAFT_84646 | Up >log2 fold | no change | no change | no annotation |
