## Supplemental figures for "Bacterial endosymbionts influence fungal transcriptional profiles with implications for host response in the human fungal pathogens *Rhizopus microsporus* and *Rhizopus delemar*": Supp-Table2.pdf

**Supplementary Table 2** Table displaying the LogFC values for a subset of genes displayed in Figure 7. (Log2 Fold Change, FDR < 0.001).

| | **Macrophage control** | **Macrophage + *R. delemar* wt** | **Macrophage + *R. microsporus* wt** | **Macrophage + *R. delemar* cured** | **Macrophage + *R. microsporus* cured** |
|---|---|---|---|---|---|
| **F3** | 2.96 | 0.195 | 2.11 | -0.593 | -2.99 |
| **PLA2G16** | 1.96 | 2.37 | 3.11 | 1.58 | -0.822 |
| **TRIM30D** | 1.96 | 2.37 | 3.11 | 1.58 | -0.822 |
| **SLC1A2** | 1.96 | 2.37 | 3.11 | 1.58 | -0.822 |
| **IL6** | 1.96 | 2.37 | 3.11 | 1.58 | -0.822 |
| **PSTPIP21** | 4.29 | 0.781 | 0.786 | -0.00761 | -2.41 |
| **SAA3** | 2.60 | 0.559 | 0.523 | -1.33 | -0.959 |
| **ENPP4** | 2.26 | 0.118 | 0.430 | -1.74 | -0.522 |
