## Supplementary figures and images for "Bacterial endosymbionts influence fungal transcriptional profiles with implications for host response in the human fungal pathogens *Rhizopus microsporus* and *Rhizopus delemar*"

### Supp-2.pdf

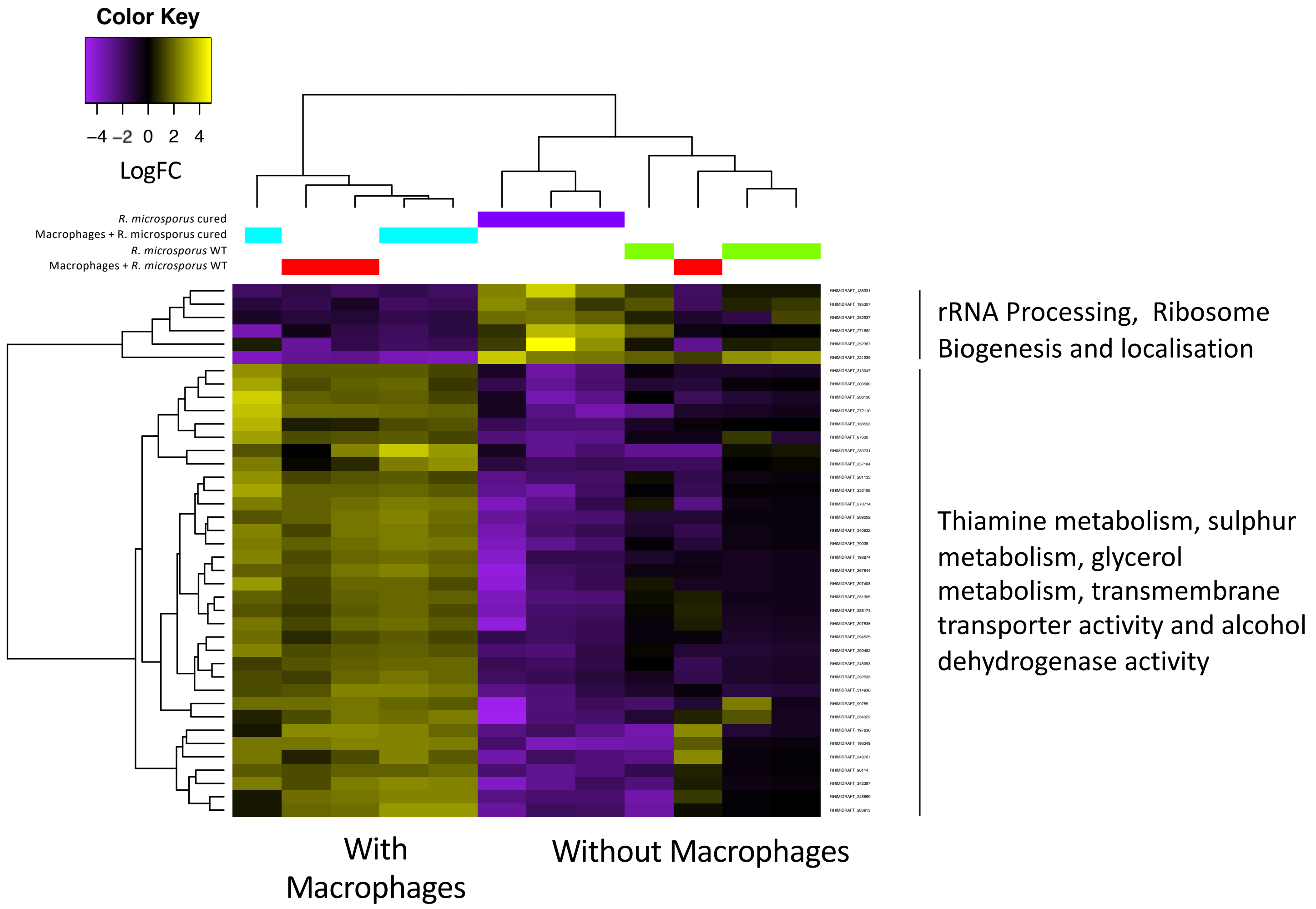

### Supp-3.pdf

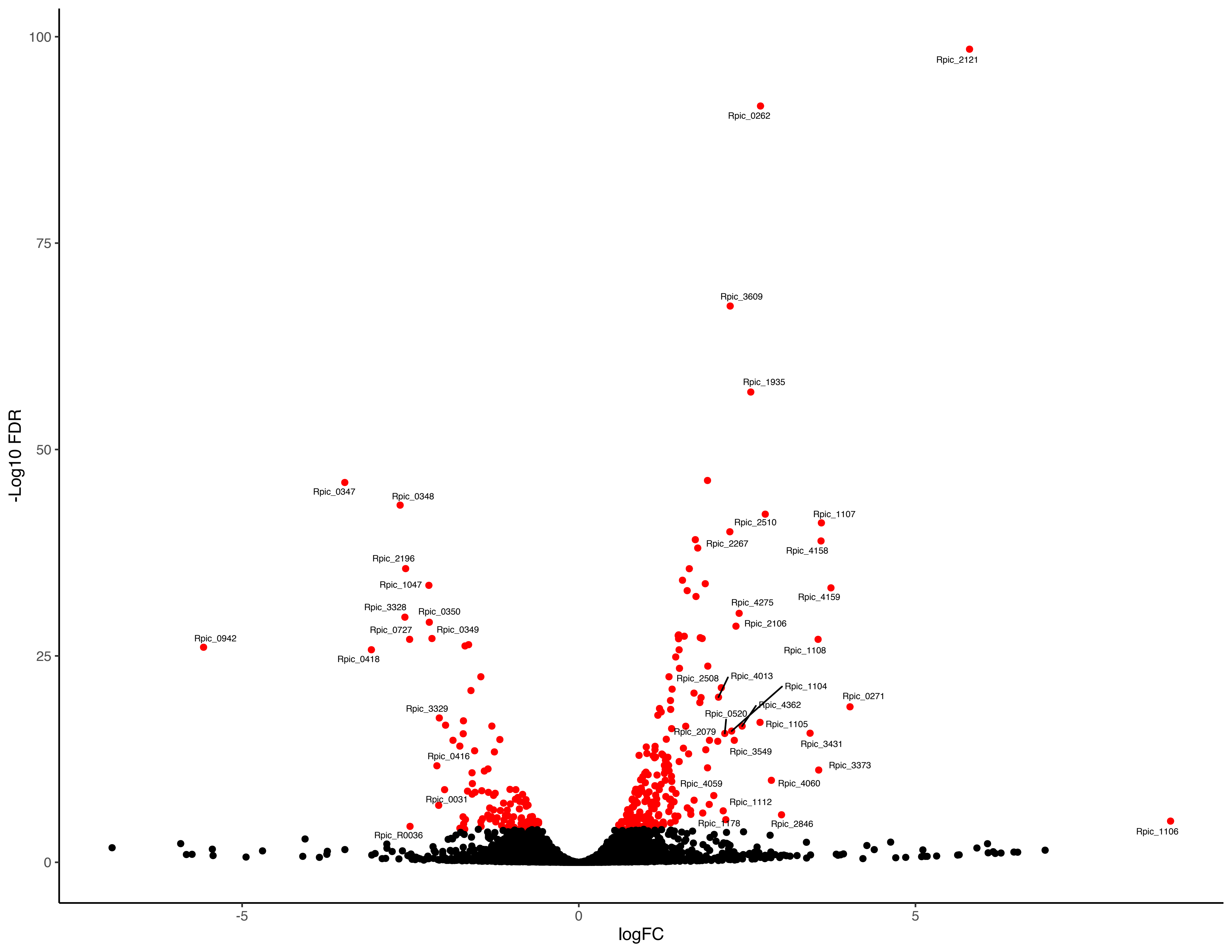

### Supp-4.pdf

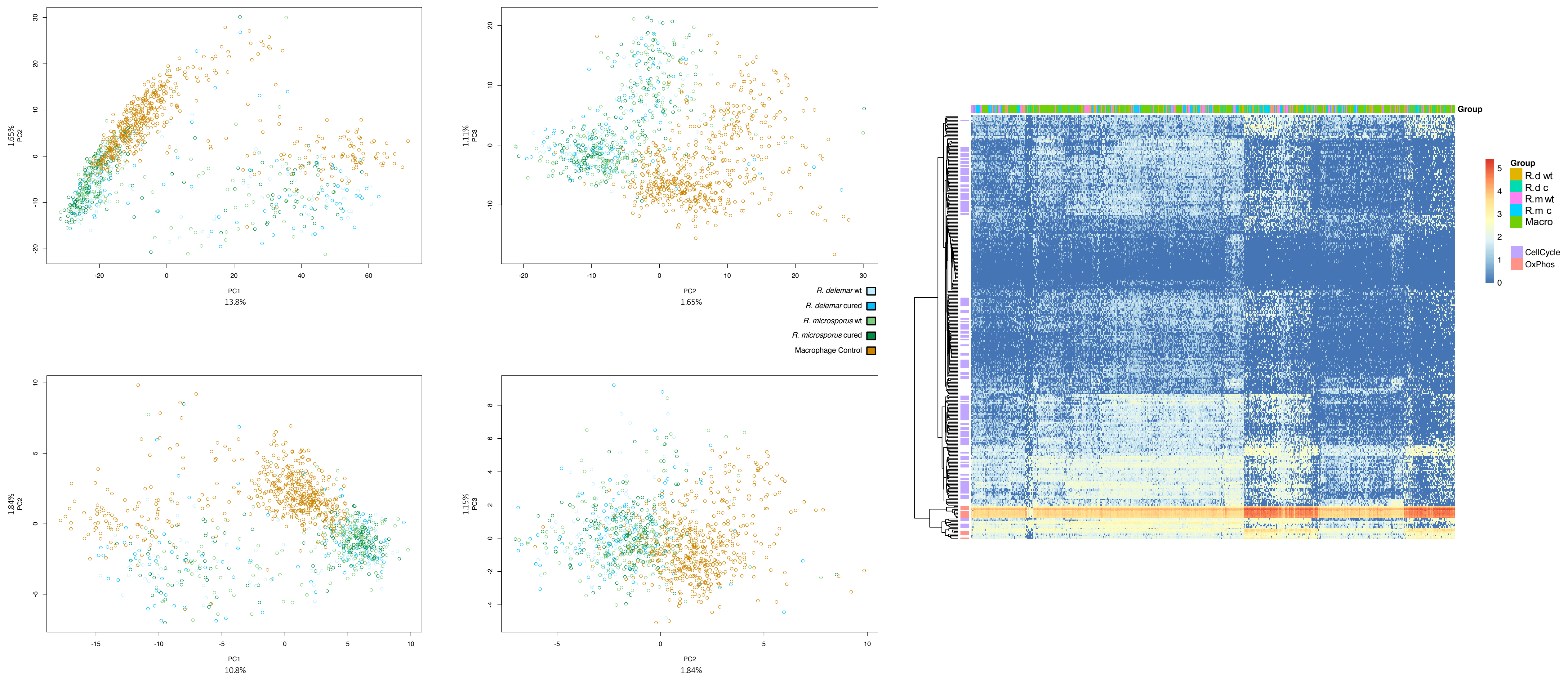

### Supp-5.pdf

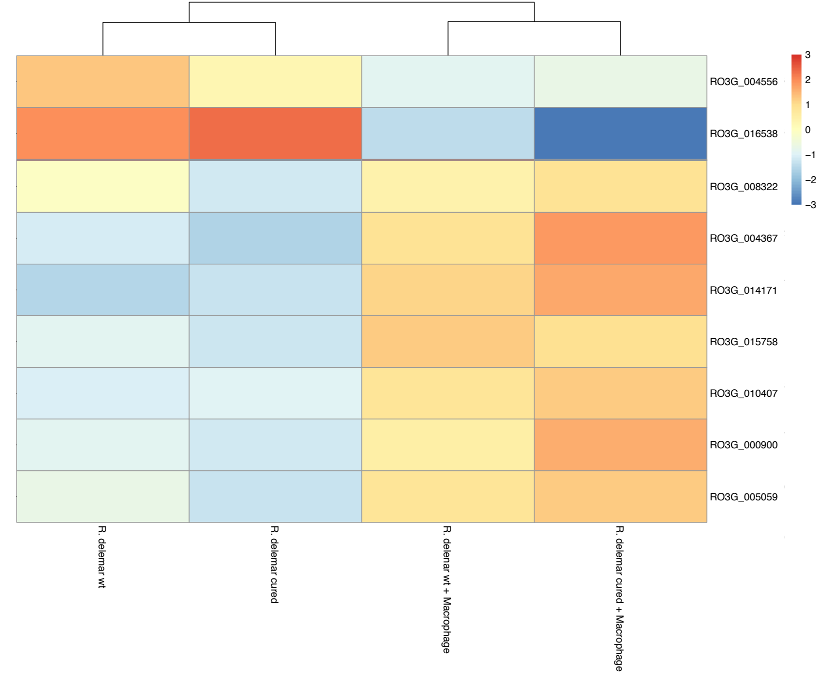

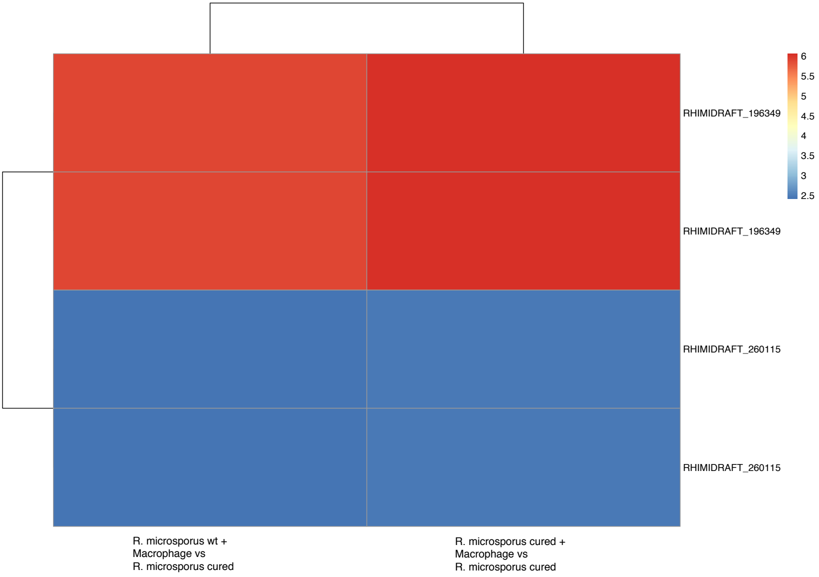

### Supp-6.pdf

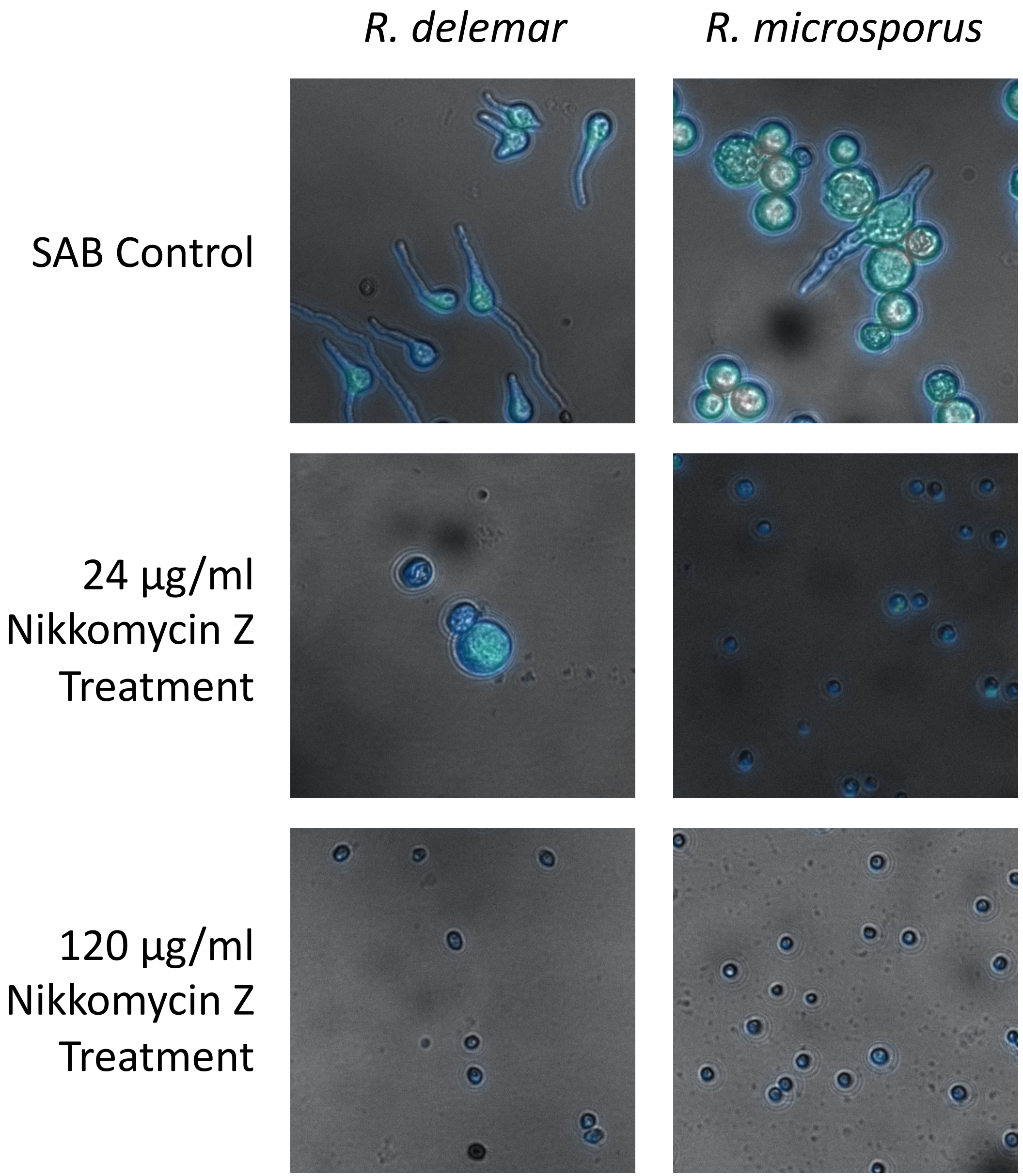
